# A PVH-to-parabrachial BDNF circuit mediates non-aversive feeding suppression

**DOI:** 10.64898/2026.09.23.753813

**Authors:** Juan Ji An, Ji-Wei Tan, Yuting Chen, Xinpei Yue, Baoji Xu

## Abstract

Parabrachial nucleus (PBN) CGRP neurons mediate aversive feeding suppression, but PBN circuitry governing homeostatic, non-aversive feeding suppression remains poorly understood. Here, we identify a circuit in which paraventricular hypothalamus (PVH) neurons innervate BDNF-expressing ventrolateral/dorsolateral PBN (vl/dlPBN^BDNF^) neurons, which project to multiple hypothalamic nuclei. Refeeding strengthens excitatory and weakens inhibitory synaptic transmission in LPBN-projecting PVH neurons and preferentially induced *Bdnf* and *Fos* in vl/dlPBN^BDNF^ neurons compared with elPBN^CGRP^ neurons, whereas noxious LPS preferentially activated elPBN^CGRP^ neurons. Gain- and loss-of-function experiments demonstrated that vl/dlPBN^BDNF^ neurons are sufficient and necessary for feeding suppression without inducing conditioned taste aversion and that this effect require BDNF-TrkB signaling. In addition to hypothalamic inputs, vl/dlPBN^BDNF^ neurons receive dense inputs from major downstream targets of elPBN^CGRP^ neurons. Thus, vl/dlPBN^BDNF^ neurons define a non-aversive feeding-suppression circuit positioned to integrate hypothalamic homeostatic signals with aversive signals, thereby coordinating food intake with internal energy needs and potentially harmful physiological conditions.

## INTRODUCTION

Food intake is regulated by two functionally distinct inhibitory processes: non-aversive and aversive feeding suppression. Non-aversive suppression reflects physiological, satiety-mediated meal termination. This process is governed by homeostatic signals, including gastric distension and hormonal feedback, acting upon neurons in the nucleus of the solitary tract (NTS) and hypothalamic nuclei like the arcuate nucleus (ARC)^1–3^. In contrast, aversive suppression is a survival-driven defense mechanism that halts consumption in response to threats such as toxins, malaise, or environmental fear^3,4^. While both processes result in the cessation of eating, they differ fundamentally in valence: satiety is a neutral or rewarding state, whereas aversive suppression involves negative affect and the formation of conditioned taste aversion (CTA)^3^.

Significant progress has been made in elucidating the neural circuits that drive aversive feeding suppression. Central to this process is a population of neurons in the external division of the lateral parabrachial nucleus that expresses calcitonin gene-related peptide (elPBN^CGRP^ neurons), which has been identified as the key mediator of aversive signals^5–9^. Activation of these neurons induces anorexia and CTA^5–7^, whereas silencing them prevents anorexia in cancer models or following the depletion of agouti-related protein (AgRP)-expressing neurons in the ARC^10,11^. The elPBN^CGRP^ neurons integrate spinal and visceral (via NTS) inputs regarding noxious stimuli^4,5,8,12^. Specific NTS populations providing direct excitatory input include noradrenergic neurons (expressing dopamine β-hydroxylase) and neurons expressing cholecystokinin (CCK)^13^. Neurons in the parabrachial nucleus (PBN) project to several downstream targets, including the capsular and lateral central amygdala (CeC and CeL), bed nucleus of the stria terminalis (BNST), gustatory thalamus, insular cortex, and lateral hypothalamus^5^. Current evidence suggests that they suppress food intake, at least in part, through their projection to the central amygdala (CeA)^5^.

Conversely, the PBN circuitry governing homeostatic, non-aversive meal termination remains poorly understood. During food consumption, vagal sensory afferents detect a variety of satiety signals produced in the gastrointestinal tract (e.g., CCK, GLP-1, and neurotransmitters) and transmit the information to the NTS^1,14–17^. Distinct groups of NTS neurons then suppress food intake by relaying these signals to regions such as the lateral parabrachial nucleus (LPBN), ARC, and paraventricular hypothalamus (PVH)^18–22^. Simultaneously, hypothalamic neurons, particularly those in the ARC, detect blood-borne adiposity and satiety signals (e.g., leptin, insulin, and GLP-1) and transmit this information to downstream control centers like the PVH^1,2^. The LPBN is a key downstream target for PVH neurons to elicit feeding suppression^23–26^. Thus, the LPBN is a converging site for the information about the states of adiposity and satiety coming through the hypothalamus and NTS, and it may play a crucial role in mediating non-aversive meal termination. However, the identity of LPBN neurons that receive inputs from the PVH remains unknown.

These LPBN neurons may mediate the anorexigenic actions of melanocortin-4 receptor (MC4R) and tropomyosin receptor kinase B (TrkB) signaling downstream of the PVH. MC4R is the receptor for alpha-melanocyte-stimulating hormone (α-MSH) and the endogenous antagonist AgRP, which are produced by proopiomelanocortin (POMC)-expressing and AgRP-expressing neurons in the ARC, respectively^2,3^. Mutations in the *MC4R* gene represent the most common monogenic cause of human obesity^27–29^. TrkB is the cognate receptor for brain-derived neurotrophic factor (BDNF)^30,31^. Extensive evidence has established BDNF-TrkB signaling as a critical regulator of energy balance^32,33^. Indeed, mutations in either the *BDNF* or the *NTRK2* (encoding TrkB) gene cause severe early-onset obesity in both mice and humans^34–42^. Notably, a case study of a child harboring a TrkB kinase domain mutation revealed caloric intake comparable to that of leptin-deficient children and greater than that observed in individuals with MC4R mutations^40^, underscoring the powerful role of this pathway. Recent studies further indicate that both MC4R and TrkB signaling regulate satiety and energy balance through PVH neurons that project to the LPBN^24,26,43^.

Here, we report that PVH neurons innervate BDNF-expressing neurons in the ventrolateral division of the PBN (vlPBN^BDNF^ neurons). Electrophysiological recordings and immunohistochemical analyses revealed that refeeding activates this pathway and enhances BDNF-TrkB signaling within it. Furthermore, chemogenetic activation of this pathway suppressed food intake without inducing CTA, whereas pharmacological inhibition of TrkB signaling with ANA-12 attenuated this anorexigenic effect. Notably, vlPBN^BDNF^ neurons are distinct not only from elPBN^CGRP^ neurons but also from previously described LPBN populations expressing prodynorphin (Pdyn) or Foxp2. Collectively, these findings identify a previously unrecognized LPBN neuronal population that serves as a postsynaptic target of PVH neurons and mediates non-aversive feeding suppression through a TrkB-dependent mechanism.

## RESULTS

### Expression of TrkB, MC4R, and GLP-1R in LPBN-projecting PVH neurons

We previously showed that deletion of the *Ntrk2* gene in LPBN-projecting PVH neurons led to hyperphagia and obesity in mice, indicating that LPBN-projecting PVH^TrkB^ neurons are involved in the control of energy balance^26^. To determine the molecular identity of these neurons, we labeled LPBN-projecting PVH neurons with mCherry by injecting retrograde AAV2retro-hSyn-mCherry into the LPBN in *Mc4r^2A-Cre/+^*;GFP-L10a/+ mice^24,44^ where GFP is expressed in MC4R-expressing cells (Figure 1A).

**Figure 1.**
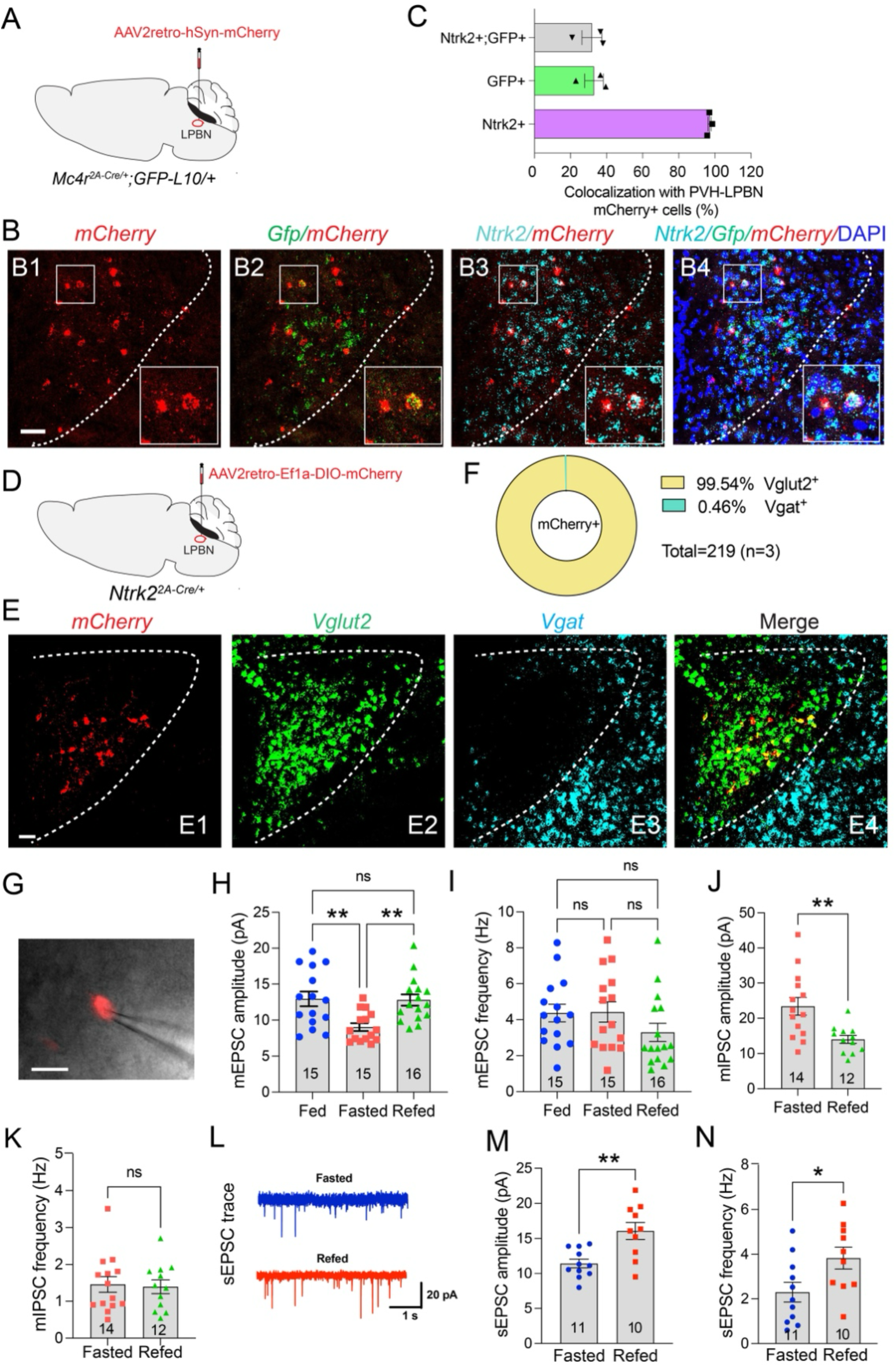
LPBN-projecting PVH^TrkB^ neurons include MC4R-expressing neurons and were activated by refeeding. (A) Strategy for labeling LPBN-projecting PVH neurons with mCherry in *Mc4r^2A-Cre/+^;GFP-L10a/+* mice where MC4R cells express GFP. (B) Representative FISH micrographs showing colocalization of *mCherry*, *GFP*, and *Ntrk2* mRNAs. Scale bar, 50 μm. (C) Quantification of mRNA colocalization. n=3 mice. (D) Strategy for labeling LPBN-projecting PVH^TrkB^ neurons with mCherry in *Ntrk2^2A-Cre/+^* mice. (E) Representative FISH micrographs showing colocalization of *mCherry*, *Slc17a6* (Vglut2), and *Slc32a1* (Vgat) mRNAs. Scale bar, 50 μm. (F) Quantification of mRNA colocalization. n=3 mice .(G) Whole-cell patch-clamp recordings of mCherry-marked LPBN-projecting PVH^TrkB^ neurons in *Ntrk2^2A-^_Cre/+_* mice. (H and I) Amplitude and frequency of mEPSCs. n=15 neurons from 4 fed mice, 15 neurons from 7 fasted mice, and 16 neurons from 7 refed mice. One-way ANOVA followed by Tukey’s multiple comparisons test: ns, not significant and \*\**p*<0.01. (J and K) Amplitude and frequency of mIPSCs. n=14 neurons from 5 fasted mice and 12 neurons from 5 refed mice. Two-tailed T test: ns, not significant and \*\**p*<0.01. (L-N) Amplitude and frequency of sEPSCs. n=11 neurons from 4 fasted mice and 10 neurons from 5 refed mice. Two-tailed T test, \**p*<0.05 and \*\**p*<0.01. All data are presented as mean ± SEM.

Fluorescence in situ hybridization (FISH) using RNAscope probes showed that nearly all LPBN-projecting PVH neurons expressed *Ntrk2* (97.1 ± 0.8%; Figures 1B and 1C) and all of them were glutamatergic as indicated by the expression of Vglut2 (Figures 1D-F). Consistent with the finding that PVH^MC4R^ neurons project to the LPBN to promote satiety^24^, approximately one third of LPBN-projecting PVH neurons were found to express GFP (indicative of MC4R expression; 33.0 ± 5.1%) or both *Ntrk2* and GFP (31.8 ± 5.5%; Figures 1B and 1C). To further validate this observation, we labeled LPBN-projecting PVH^MC4R^ neurons with mCherry (Figure S1A), conducted FISH with probes for *mCherry* and *Ntrk2* mRNAs (Figure S1B), and found that 84% of these neurons expressed *Ntrk2* (Figure S1C).

GLP-1R-expressing PVH neurons represent another neuronal population known to suppress feeding^25,45^. These neurons accounted for approximately one quarter of LPBN-projecting PVH neurons (25.2 ± 1.4%), and about one third of them also expressed *Mc4r* mRNA (Figures S1D-F).

Collectively, the FISH analyses indicate that glutamatergic PVH^TrkB^ neurons comprise nearly all LPBN-projecting PVH neurons, including the vast majority of LPBN-projecting PVH^MC4R^ and PVH^GLP-1R^ neurons. These findings suggest that LPBN neurons receiving excitatory inputs from PVH^TrkB^ neurons mediate the anorectic effects not only of TrkB signaling but also of MC4R and GLP-1R signaling.

### Refeeding activates LPBN-projecting PVH^TrkB^ neurons

We previously found that 2-hour refeeding after an overnight fast increased Fos expression in LPBN-projecting PVH^TrkB^ neurons^26^, suggesting that refeeding activates these neurons. To examine their synaptic transmission, we labeled LPBN-projecting PVH^TrkB^ neurons with mCherry by injecting AAV2retro-Ef1a-DIO-mCherry into the LPBN of adult *Ntrk2^2A-Cre/+^* mice (Figure 1D). Whole-cell patch-clamp recordings were then performed in mCherry-labeled neurons (Figure 1G) to measure miniature excitatory postsynaptic currents (mEPSCs) and miniature inhibitory postsynaptic currents (mIPSCs) in the presence of tetrodotoxin (TTX), which blocks action potential-dependent neurotransmitter release. In a separate set of recordings, spontaneous excitatory postsynaptic currents (sEPSCs) were measured in the absence of TTX.

We found that 2-hour refeeding increased the amplitude, but not the frequency, of mEPSCs in LPBN-projecting PVH^TrkB^ neurons (Figures 1H and 1I). These findings suggest that refeeding enhances excitatory postsynaptic strength without substantially altering basal presynaptic neurotransmitter release. Because mEPSC amplitude and frequency were comparable between fed and refed mice, subsequent recordings focused on the fasted and refed states. Refeeding also decreased the amplitude of mIPSCs without altering their frequency (Figures 1J and 1K), indicating a reduction in inhibitory postsynaptic strength, potentially through changes in the number or conductance of GABA_A_ receptors. Interestingly, refeeding increased both the amplitude and frequency of sEPSCs in LPBN-projecting PVH^TrkB^ neurons (Figures 1M and 1N). The increase in sEPSC amplitude is consistent with the enhancement of excitatory postsynaptic strength observed in mEPSC recordings, whereas the increase in sEPSC frequency indicates elevated activity of presynaptic excitatory neurons and/or increased action potential-dependent glutamate release. Collectively, these results suggest that refeeding activates LPBN-projecting PVH^TrkB^ neurons through multiple mechanisms, including enhanced excitatory synaptic strength, reduced inhibitory synaptic strength, and increased excitatory input from upstream neurons.

### LPBN-projecting PVH^TrkB^ neurons suppress food intake in a TrkB signaling-dependent manner

We used designer receptors exclusively activated by designer drugs (DREADDs) to determine whether LPBN-projecting PVH^TrkB^ neurons regulate food intake. To selectively target these neurons, we stereotaxically injected retrograde AAV2retro-Ef1a-DIO-FLPo into the LPBN and either FLPo-dependent AAV2-Ef1a-fDIO-mCherry or AAV5-hSyn-fDIO-hM3D(Gq)-mCherry into the PVH of adult *Ntrk2^2A-Cre/+^* mice (Figure 2A). Retrograde transport of AAV2retro-Ef1a-DIO-FLPo to the PVH induced FLPo expression specifically in LPBN-projecting PVH^TrkB^ neurons, thereby driving selective expression of either mCherry or the excitatory DREADD hM3D(Gq)-mCherry (Figure 2B).

**Figure 2.**
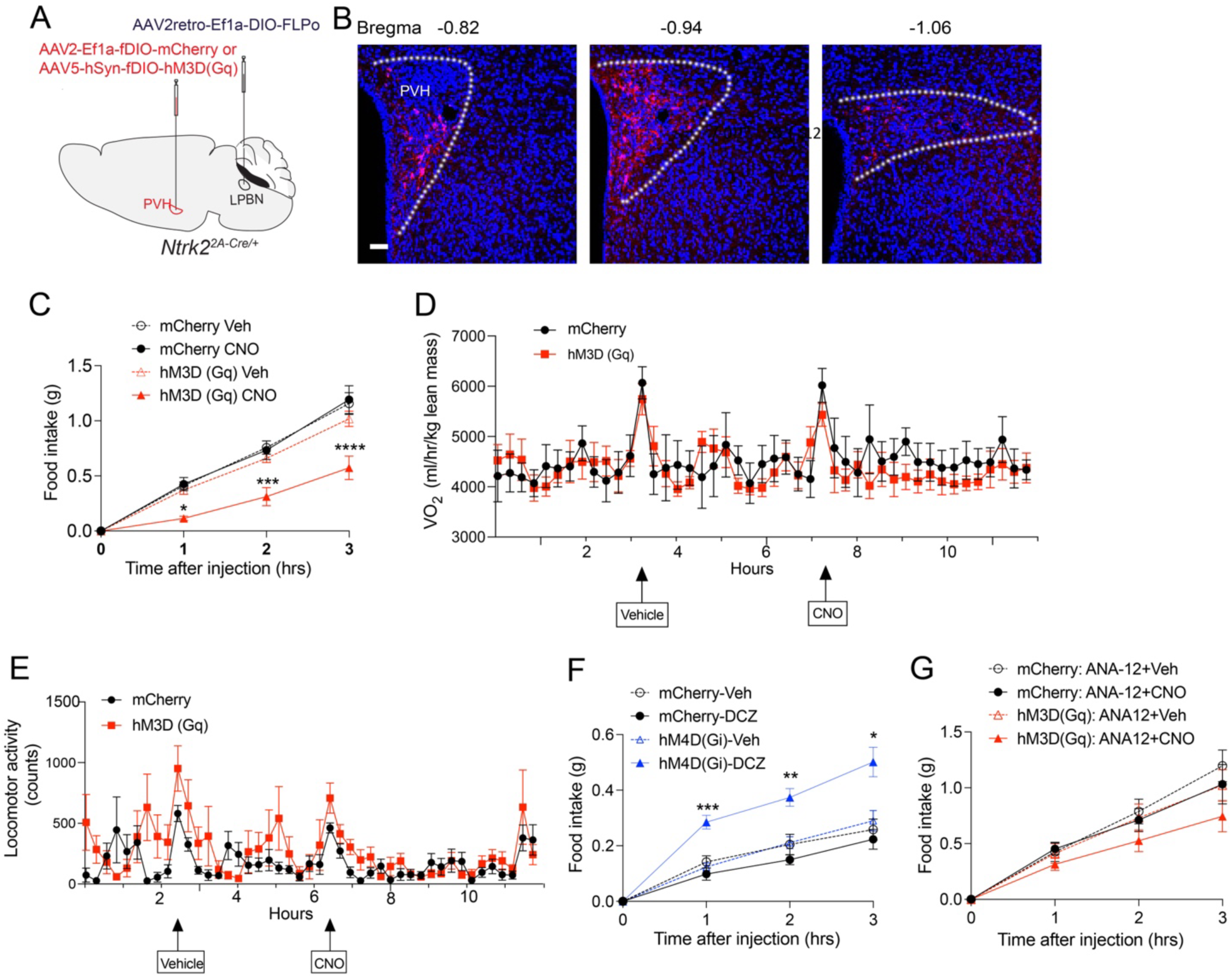
Activation of LPBN-projecting PVH^TrkB^ neurons suppresses food intake in a TrkB signaling-dependent manner. (A) Strategy for selective expression of mCherry or hM3D(Gq)-mCherry in LPBN-projecting PVH^TrkB^ neurons. (B) Confocal micrographs showing mCherry expression in PVH neurons across the rostrocaudal axis. Scale bar, 50 μm. (C) Chemogenetic activation of LPBN-projecting PVH^TrkB^ neurons suppresses nocturnal food intake. n=7 mice for mCherry group and 8 mice for hM3D(Gq) group. Two-way ANOVA followed by Bonferroni’s multiple comparisons test, *p<0.05, ***p<0.001 and ****p< 0.0001. (D) Oxygen consumption (VO_2_). n=6-8 mice per group. Two-way ANOVA followed by Šídák’s multiple comparisons test, not significant. (E) Locomotor activity measured by x-axis beam breaks. n=7 mice for mCherry group and 8 mice for hM3D(Gq) group. Two-way ANOVA followed by Sidak’s multiple comparison test, not significant. (F) Chemogenetic inhibition of LPBN-projecting PVH^TrkB^ neurons increases daytime food intake. n=6 mice per group. Two-way ANOVA followed by Bonferroni’s multiple comparisons test, *p<0.05, **p<0.01 and ***p< 0.001. (G) ANA-12 pretreatment abolishes the suppressive effect of LPBN-projecting PVH^TrkB^ neuron activation on nocturnal food intake. n=7 mice for mCherry group and 8 mice for hM3D(Gq) group. Two-way ANOVA followed by Bonferroni’s multiple comparisons test, not significant. All data are presented as mean ± SEM.

Administration of clozapine N-oxide (CNO; 1 mg/kg, i.p.) significantly suppressed nocturnal food intake in mice expressing hM3D(Gq) without affecting oxygen consumption or locomotor activity (Figures 2C-2E). To determine whether LPBN-projecting PVH^TrkB^ neurons are required for satiety, we selectively expressed the inhibitory DREADD hM4D(Gi) in these neurons. Administration of deschloroclozapine (DCZ), a potent hM4D(Gi) agonist^46^, increased daytime food intake, when mice are normally inactive (Figure 2F). Together, these gain- and loss-of-function experiments demonstrate that the PVH^TrkB^-to-LPBN circuit is both sufficient and necessary to promote satiety while having minimal effects on energy expenditure and locomotor activity.

BDNF enhances glutamatergic synaptic transmission while weakening GABAergic transmission by modulating activity-dependent neurotransmitter release and postsynaptic responsiveness^47^. Because refeeding increased excitatory input and decreased inhibitory input onto LPBN-projecting PVH^TrkB^ neurons (Figures 1G-1N), we hypothesized that refeeding activates these neurons through BDNF-TrkB signaling. If this model is correct, pharmacological inhibition of TrkB signaling should attenuate the anorexigenic effect of activating these neurons. Consistent with this hypothesis, pretreatment with the selective TrkB antagonist ANA-12^48^ significantly reduced the suppression of nocturnal food intake induced by CNO administration in mice expressing hM3D(Gq) in LPBN-projecting PVH^TrkB^ neurons (Figure 2G). These findings indicate that TrkB signaling is required for the full feeding-suppressive effect of activating the PVH^TrkB^-to-LPBN circuit, supporting a role for BDNF-TrkB signaling in regulating the activity of this pathway.

### BDNF neurons in the vlPBN respond to refeeding

Because BDNF is the endogenous ligand for TrkB and nearly all LPBN-projecting PVH neurons express TrkB (Figure 1C), we hypothesized that their postsynaptic targets in the LPBN express BDNF. To identify these postsynaptic neurons, we took advantage of the anterograde transsynaptic properties of AAV1^49^. We injected AAV1-Ef1a-DIO-FLPo into the PVH and FLPo-dependent AAV2-Ef1a-fDIO-mCherry into the LPBN of *Bdnf^2A-Cre/+^*mice, enabling selective labeling of BDNF-expressing neurons that receive input from PVH^TrkB^ neurons (Figure 3A). mCherry-labeled neurons were concentrated in the vlPBN, with fewer labeled neurons in the dorsolateral PBN (dlPBN) and elPBN (Figure 3B). For simplicity, we hereafter refer to BDNF-expressing neurons in the vlPBN and dlPBN collectively as vl/dlPBN^BDNF^ neurons.

**Figure 3.**
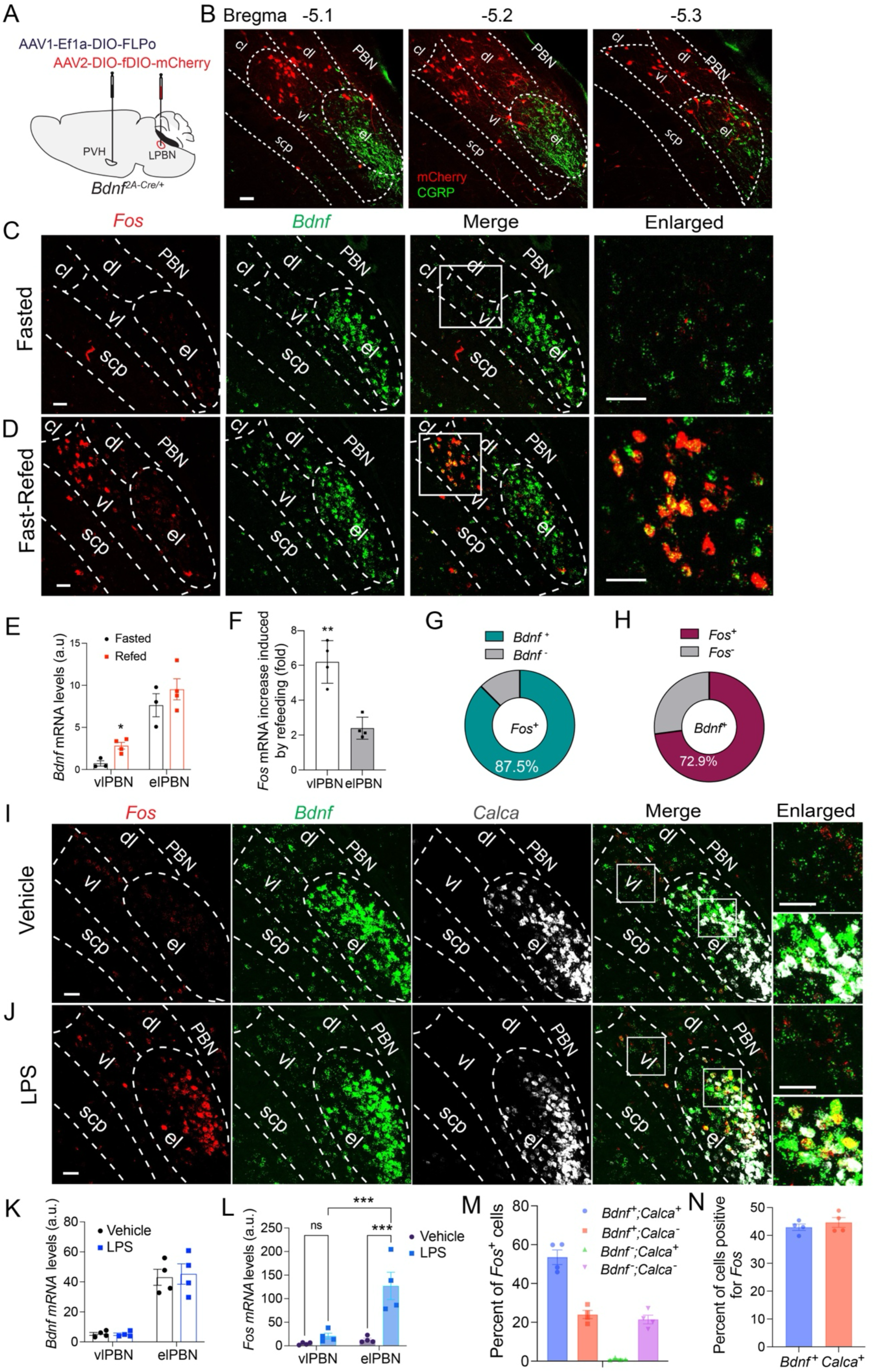
vlPBN^BDNF^ neurons are postsynaptic to PVH^TrkB^ neurons and respond to refeeding. (A) Strategy for labeling BDNF-expressing LPBN (LPBN^BDNF^) neurons postsynaptic to PVH neurons in *Bdnf^2A-Cre/+^* mice. (B) Distribution of LPBN^BDNF^ neurons that are postsynaptic targets of PVH neurons. CGRP immunohistochemistry (green) marks the elPBN. (C, D) Representative FISH images showing *Bdnf* and *Fos* mRNA expression in mice fasted overnight or refed for 2-h following an overnight fast. The enlarged images in the rightmost column correspond to the boxed regions in the third column and highlight the expression of *Bdnf* and *Fos* mRNAs in the vlPBN. (E) *Bdnf* mRNA levels in the vlPBN and elPBN of fasted and refed mice. Unpaired Student’s t test, *p<0.05. n=3-4 mice. (F) *Fos* mRNA levels in refed mice relative to fasted mice. Unpaired Student’s t test, **p<0.01, n=4 mice. (G, H) Colocalization of *Bdnf* and *Fos* mRNAs in vlPBN of refed mice (n=4 mice). (I, J) Representative FISH images showing *Bdnf* and *Fos* mRNA expression in mice treated with either vehicle or LPS. Brain tissue was collected 90 min after LPS injection. The enlarged images in the rightmost column correspond to the boxed regions in the fourth column and highlight the expression of *Bdnf*, *Fos*, and *Calca* mRNAs in the elPBN. (K) *Bdnf* mRNA levels in the vlPBN and elPBN of vehicle- or LPS-treated mice. Unpaired Student’s t test, not significant. n=4 mice. (L) *Fos* mRNA levels in the vlPBN and elPBN of LPS-treated mice relative to vehicle-treated mice. Unpaired Student’s t test, ***p<0.001, n=4 mice. (M, N) Colocalization of *Bdnf*, *Calca*, and *Fos* mRNAs in the elPBN of LPS-treated mice (n=4 mice). All data are presented as mean ± SEM. Scale bars, 50 μm. vl, ventral lateral PBN; el, external lateral PBN.

Because AAV1 exhibits limited retrograde transport^49^, we used mono-synaptic retrograde tracing with a modified rabies virus system^50–54^ to independently validate the PVH^TrkB^-to-vl/dlPBN^BDNF^ synaptic connection. The EnvA-pseudotyped glycoprotein-deleted rabies virus, SADΔG-EGFP(EnvA), requires expression of the avian tumor virus receptor A (TVA) and rabies glycoprotein (RG) for selective infection and monosynaptic retrograde spread^53^. To target LPBN^BDNF^ neurons, we injected AAV2-CMV-FLEX-TVAmCherry-2A-oG^55,56^ into the LPBN of *Bdnf^2A-Cre/+^* mice, thereby inducing expression of TVA-mCherry and optimized RG (oG) in BDNF-expressing neurons (Figures S2A and S2B). Three weeks later, SADΔG-EGFP(EnvA) was injected into the LPBN, where it selectively infected TVA-expressing LPBN^BDNF^ neurons and spread retrogradely to their direct presynaptic partners in an oG-dependent manner^57,58^. Consistent with our hypothesis, EGFP-labeled presynaptic neurons were detected in the PVH and expressed *Ntrk2* mRNA, confirming a direct synaptic connection between PVH^TrkB^ neurons and LPBN^BDNF^ neurons (Figure S2C).

To investigate the role of vl/dlPBN^BDNF^ neurons in feeding regulation, we examined BDNF and Fos expression in response to refeeding, with a particular emphasis on the vlPBN, where most PVH-innervated BDNF neurons are located. *Bdnf* mRNA was expressed at higher levels in the elPBN than in the vlPBN under both fasted and refed conditions (Figures 3C and 3D). In the elPBN, *Bdnf* expression largely overlapped with *Calca* mRNA, which encodes CGRP, although a subset of neurons expressed *Bdnf* without *Calca* (Figure S2E). Despite the abundance of BDNF neurons in the elPBN, only a few received input from PVH^TrkB^ neurons (Figure 3B). Notably, 2-hour refeeding after an overnight fast significantly increased *Bdnf* mRNA expression in the vlPBN but not elPBN (Figures 3C-3E). Refeeding also induced *Fos* mRNA more robustly in the vlPBN than in the elPBN (Figures 3C, 3D, 3F, and S2F), as previously reported^7^. Within the vlPBN, 88% of Fos^+^ cells expressed *Bdnf*, whereas 73% of BDNF neurons expressed Fos following refeeding (Figures 3G and 3H), indicating that vlPBN^BDNF^ neurons are strongly recruited by physiological satiety signals. In contrast, intraperitoneal LPS administration, which mimics bacterial infection and produces sickness behavior including anorexia, did not significantly increase the expression of either *Bdnf* or *Fos* in vl/dlPBN neurons (Figures 3I-3L and S2G), while strongly stimulating *Fos* expression in the elPBN as expected (Figures 3I-3N and S2G).

Because elPBN^CGRP^ neurons suppress feeding in response to visceral malaise and external threats through aversive mechanisms^7,8,11^, these findings suggest that vl/dlPBN^BDNF^ neurons, rather than elPBN^CGRP^ neurons, play the predominant role in the homeostatic regulation of food intake.

### LPBN^BDNF^ neurons postsynaptic to the PVH mediate feeding suppression

To directly determine the role of LPBN^BDNF^ neurons postsynaptic to the PVH in the homeostatic regulation of food intake, we selectively expressed either mCherry (control) or the excitatory DREADD hM3D(Gq) in these neurons. Anterograde AAV1-Ef1a-DIO-FLPo virus was injected into the PVH, and either AAV2-Ef1a-fDIO-mCherry or AAV5-hSyn-fDIO-hM3D(Gq)-mCherry was injected into the LPBN of 8-week-old *Bdnf^2A-Cre/+^* mice (Figures 4A-4C). AAV2-CMV-EGFP was co-injected into the PVH to label the injection site (Figure 4B). hM3D(Gq)-expressing neurons were located predominantly in the vlPBN, with smaller populations in the dlPBN and elPBN (Figure 4C). Administration of CNO markedly suppressed nocturnal food intake in mice expressing hM3D(Gq), but not in control mice (Figure 4D). These results indicate that chemogenetic activation of PVH-innervated LPBN^BDNF^ neurons is sufficient to suppress feeding.

**Figure 4.**
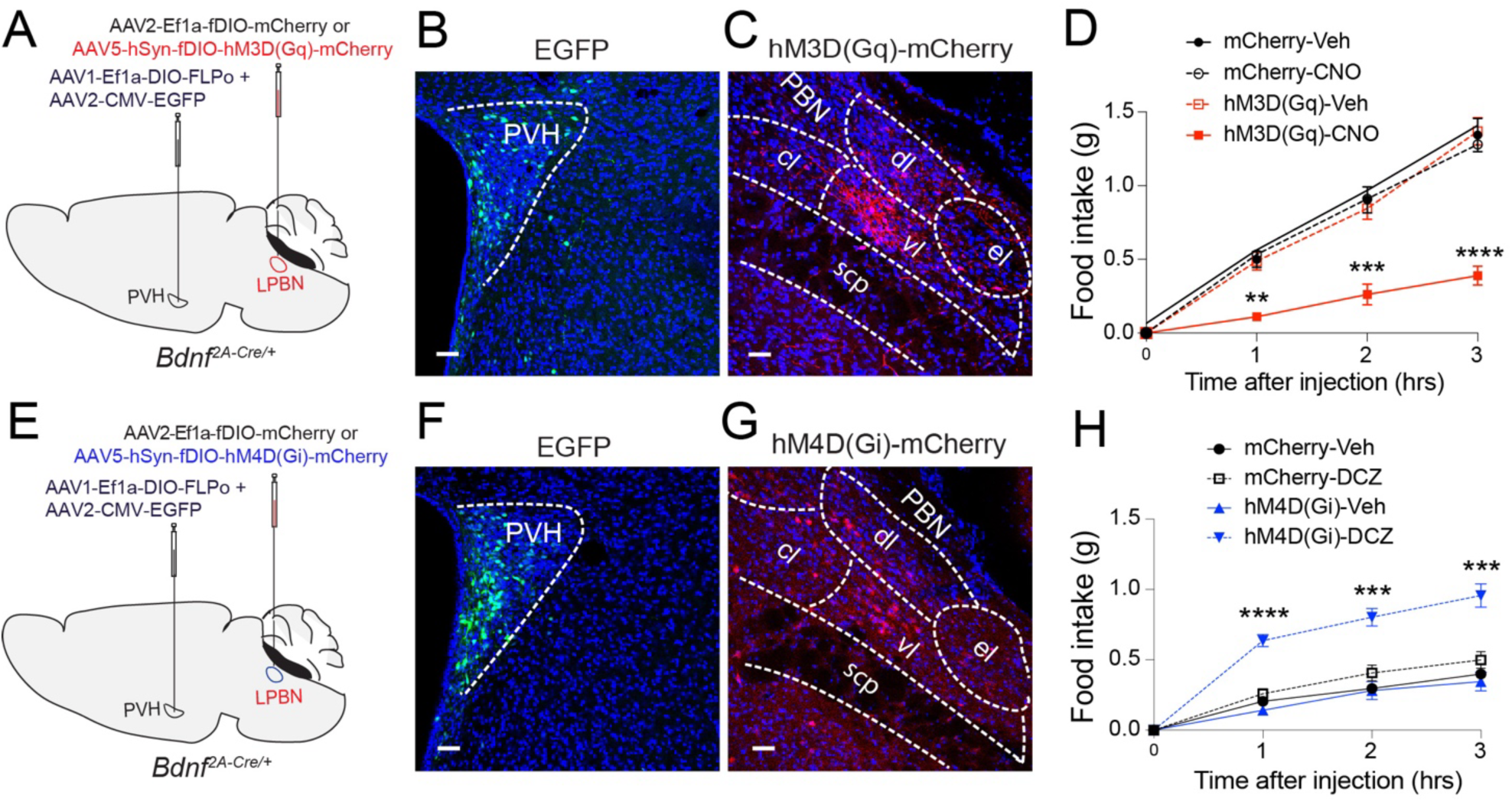
LPBN^BDNF^ neurons postsynaptic to the PVH regulate food intake. (A) Strategy for selective expression of either mCherry or hM3D(Gq)-mCherry in PVH-innervated LPBN^BDNF^ neurons. (B) EGFP expressed from co-injected AAV2-CMV-EGFP marked the injection site in the PVH. Scale bar, 50 μm. (C) hM3D(Gq)-mCherry expression in the LPBN. Scale bar, 50 μm. (D) Nocturnal food intake following vehicle (Veh) or CNO injection. n=8 mice per group. Two-way ANOVA followed by Tukey’s multiple comparisons test, ***p<0.001 and ****p< 0.0001. (E) Strategy for selective expression of either mCherry or hM4D(Gi)-mCherry in PVH-innervated LPBN^BDNF^ neurons. (F) EGFP expressed from co-injected AAV2-CMV-EGFP marked the injection site in the PVH. Scale bar, 50 μm. (G) hM4D(Gi)-mCherry expression in the LPBN. Scale bar, 50 μm. (H) Food intake during the light cycle following vehicle (Veh) or DCZ injection. n=6 mice per group. Two-way ANOVA followed by Tukey’s multiple comparisons test, ***p<0.001 and ****p< 0.0001. All data are presented as mean ± SEM.

Using the same viral strategy, we selectively expressed either mCherry (control) or the inhibitory DREADD hM4D(Gi) in these neurons (Figures 4E-4G). Administration of DCZ robustly increased light-cycle food intake in mice expressing hM4D(Gi), but not in control mice (Figure 4H), indicating the activity of these neurons is required for normal satiety.

Collectively, these gain- and loss-of-function experiments demonstrate that LPBN^BDNF^ neurons postsynaptic to the PVH, located predominantly in the vlPBN, are both sufficient and necessary for feeding suppression.

### Vl/dlPBN^BDNF^ neurons constitute a distinct neuronal population

Two neuronal populations in the LPBN have previously been implicated in feeding regulation. Palatable food activates *Foxp2*-expressing neurons in the LPBN, including the vlPBN, whereas optogenetic activation of prodynorphin (Pdyn)-expressing neurons in the PBN induces aversive feeding suppression^59,60^. To determine the relationship between these neuronal populations and BDNF-expressing neurons in the vlPBN, we examined co-expression of *Bdnf*, *Foxp2*, and *Pdyn* across the rostrocaudal axis (Figures 5A and 5B).

**Figure 5.**
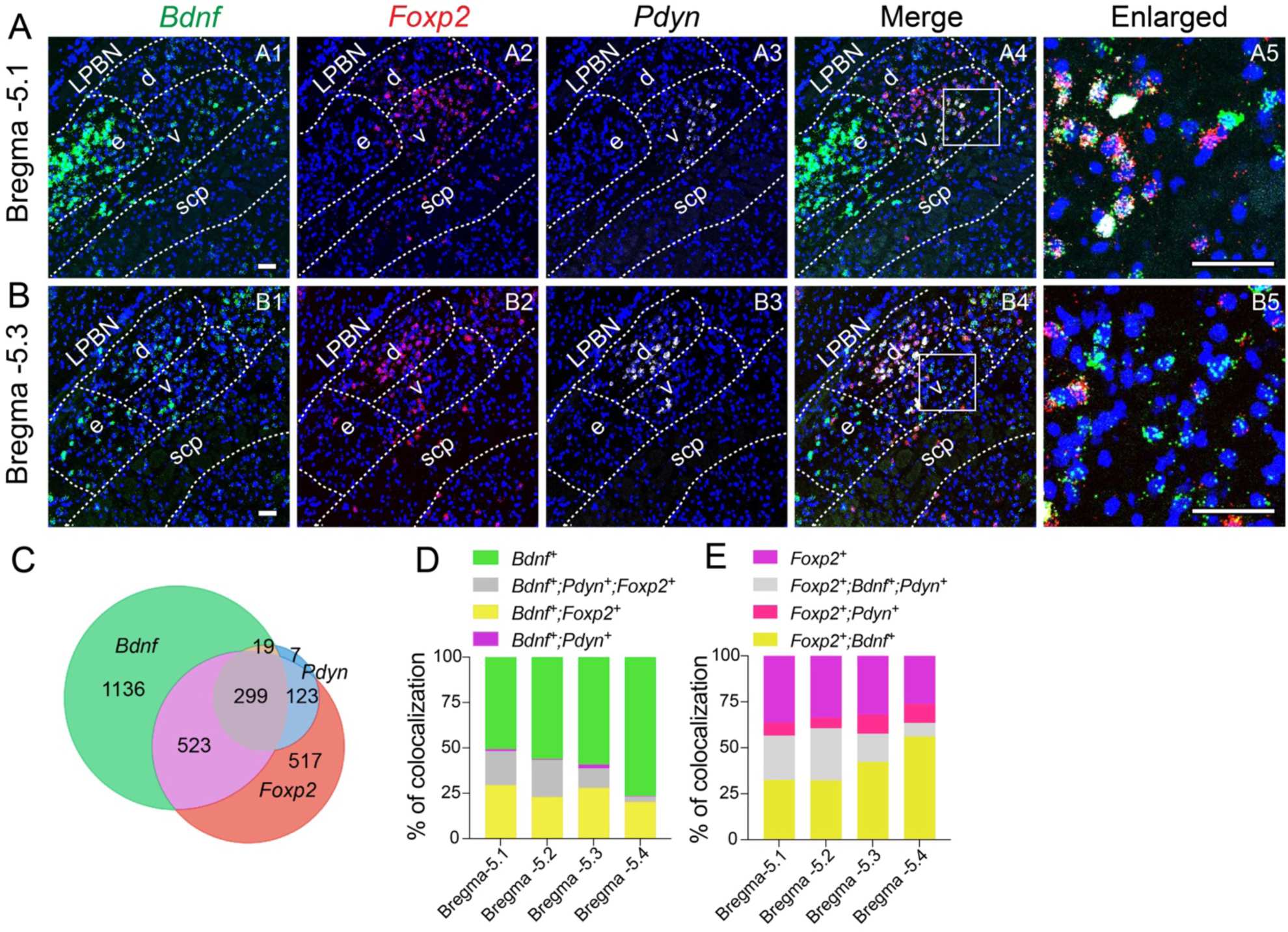
Expression of *Bdnf*, *Foxp2*, and *Pdyn* mRNAs in the vlPBN. (A, B) Representative FISH images showing the distribution of *Bdnf*, *Foxp2*, and *Pdyn* mRNAs at two positions along the rostrocaudal axis of the vlPBN. A5 and B5 show higher-magnification views of the boxed areas in A4 and B4, respectively. Abbreviations: d, dlPBN; e, elPBN; v, vlPBN. Scale bars, 50 μm. (C) Venn diagram showing vlPBN neurons co-expressing *Bdnf*, *Foxp2*, and *Pdyn* mRNAs. n=4 mice. (D) Fractions of vlPBN^BDNF^ neurons that co-express *Foxp2* mRNA (yellow), *Pdyn* mRNA (purple), both transcripts (gray), or neither transcript (green) at four rostrocaudal levels of the vlPBN. n=4 mice. (E) Fractions of vlPBN^Foxp2^ neurons that co-express *Bdnf* mRNA (yellow), *Pdyn* mRNA (pink), both transcripts (gray), or neither transcript (purple) at four rostrocaudal levels of the vlPBN. n=4 mice.

*Bdnf* mRNA was broadly expressed throughout the LPBN, with the highest expression in the elPBN (Figures 5A1 and 5B1). In contrast, *Foxp2* and *Pdyn* mRNAs were largely restricted to the vlPBN and dlPBN, with minimal expression in the elPBN (Figures 5A2, 5A3, 5B2, and 5B3). Within the vlPBN, *Foxp2* and *Pdyn* expression was concentrated in the rostral region (Bregma -5.1 mm), whereas *Bdnf* was distributed throughout the rostral and caudal regions (Figures 5A1-5A3 and 5B1-5B3). Although substantial overlap was observed between *Bdnf* mRNA and either *Foxp2* or *Pdyn* mRNA, particularly in the rostral vlPBN (Figures 5A4, 5A5, 5B4, and 5B5), their overall distribution differed.

Quantitative analysis revealed that BDNF-expressing neurons were the most abundant neuronal population in the vlPBN, whereas *Pdyn*-expressing neurons were the least abundant (Figure 5C). Nearly all *Pdyn*-expressing neurons co-expressed *Foxp2*, whereas more than half of vlPBN^BDNF^ neurons lacked either *Foxp2* or *Pdyn* expression, particularly in the caudal vlPBN (Figures 5C and 5D). Conversely, more than half of vlPBN^Foxp^^2^ neurons expressed *Bdnf* (Figure 5E). These findings demonstrate that the vl/dlPBN^BDNF^ neurons comprise a molecularly distinct neuronal population that partially overlaps with, but is substantially broader than, the *Foxp2*- and *Pdyn*-expressing populations. This molecular distinction prompted us to determine whether *Foxp2*- and *Pdyn*-expressing neurons, like vl/dlPBN^BDNF^ neurons, contribute to feeding suppression.

### Activating PVH-innervated LPBN^Foxp^^2^ or LPBN^Pdyn^ neurons is insufficient to suppress food intake

The gene expression analyses described above indicate that vlPBN neurons can be subdivided into distinct populations based on their expression and co-expression of *Bdnf*, *Foxp2,* and *Pdyn* (Figure 5). To determine which populations are activated by physiological satiety signals, we examined *Fos* mRNA induction after 2 h of refeeding following an overnight fast (Figures S3A and S4A).

Refeeding induced robust *Fos* expression throughout the vlPBN (Figures S3A1, S3A5, S4B1, and S4B3). Most *Fos*^+^ neurons expressed *Bdnf* (83.2 ± 1.0%; n=4 mice), and 44% of refeeding-activated *Bdnf* neurons also expressed *Foxp2* (Figure S3B). The proportion of *Fos*^+^ neurons expressing *Bdnf* remained relatively constant along the rostrocaudal axis, whereas the proportion expressing *Foxp2* decreased caudally (Figure S3C), consistent with the more restricted distribution of *Foxp2*-expressing neurons (Figure 5). Because *Pdyn*-expressing neurons comprise only a small fraction of vlPBN neurons, relatively few *Fos*^+^ vlPBN neurons expressed *Pdyn*, particularly in caudal sections (Figures S4C and S4D).

Interestingly, similar proportions of the *Bdnf*-, *Foxp2*-, and *Pdyn*-expressing populations were activated by refeeding (Figures 3H, S3D, and S4E). However, because BDNF-expressing neurons are considerably more abundant than either *Foxp2*- or *Pdyn*-expressing neurons (Figure 5C), they constitute the majority of refeeding-responsive neurons in the vlPBN.

To determine whether *Foxp2*- or *Pdyn*-expressing neurons mediate feeding suppression, we selectively expressed either mCherry or hM3D(Gq) in PVH-innervated LPBN^Foxp2^ or LPBN^Pdyn^ neurons using *Foxp2^Cre/+^* or *Pdyn^IRES-Cre/+^* mice (Figures S5A, S5B, S5D, and S5E). Unlike activation of PVH-innervated LPBN^BDNF^ neurons (Figure 4), chemogenetic activation of either LPBN^Foxp2^ or LPBN^Pdyn^ neurons failed to suppress nocturnal food intake (Figures S5C and S5F). These findings indicate that neither the *Foxp2*-nor the *Pdyn*-expressing population alone is sufficient to mediate feeding suppression, suggesting that the relevant satiety-promoting neurons are contained within the broader LPBN^BDNF^ population.

### Distinct projections of elPBN^CGRP^ and PVH-innervated LPBN^BDNF^ neurons

We next sought additional anatomical evidence that elPBN^CGRP^ and PVH-innervated LPBN^BDNF^ neurons play distinct roles in feeding regulation by comparing their axonal projections. To trace elPBN^CGRP^ projections, we injected AAV2-CAG-DIO-tdTomato into the LPBN of *Calca^Cre/+^* mice (Figures S6A and S6B). Consistent with previous findings^5^, elPBN^CGRP^ neurons projected to the bed nucleus of the stria terminalis (BNST), the capsular and lateral part of the central amygdala (CeC and CeL), the parvicellular part of the ventral posteromedial thalamic nucleus (VPMpc), and the parasubthalamic nucleus (PSTN), but showed little or no innervation of the hypothalamus, including the PVH, DMH, and ARC (Figures S6C1-S6C7).

To map the projections of PVH-innervated LPBN^BDNF^ neurons, we injected AAV1-Ef1a-DIO-FLPo together with AAV-GFP into the PVH and FLPo-dependent AAV2-Ef1a-fDIO-mCherry into the LPBN of *Bdnf^2A-Cre/+^* mice (Figures 6A and 6B). As described above, labeled neurons were located predominantly in the vlPBN, with smaller populations in the dlPBN and elPBN (Figures 6C-6E). In contrast to elPBN^CGRP^ neurons, PVH-innervated LPBN^BDNF^ neurons projected extensively to the hypothalamic regions, including the PVH, DMH, and ARC, with only sparse projections to the CeA and BNST (Figures 6F-6M).

**Figure 6.**
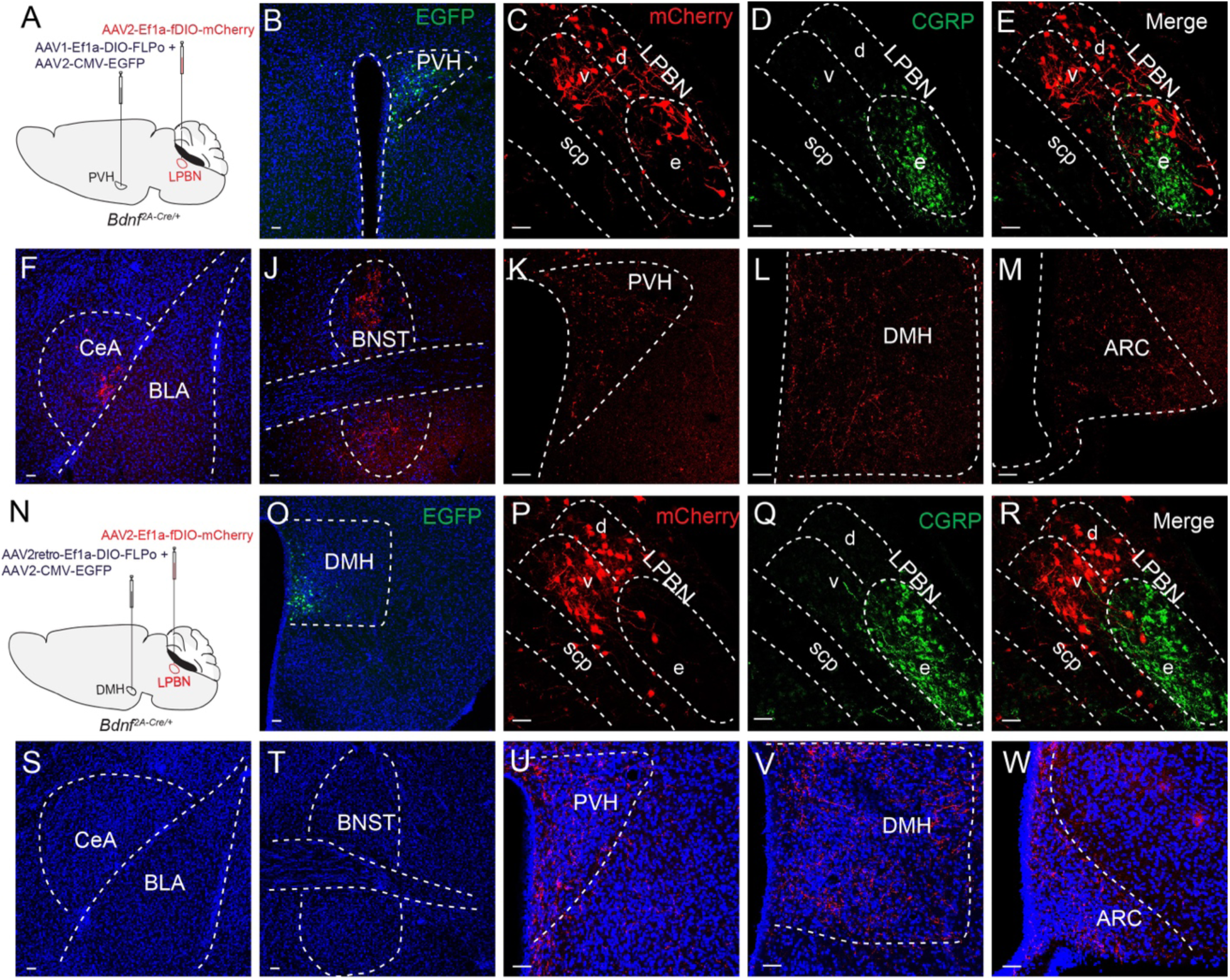
Projections of PVH-innervated LPBN^BDNF^ neurons. (A) Strategy for selective labeling of PVH-innervated LPBN^BDNF^ neurons in *Bdnf^2A-Cre/+^* mice using anterograde AAV1. (B) EGFP marks the injection site in the PVH. Scale bar, 50 μm. (C-E) mCherry-labeled PVH-innervated LPBN^BDNF^ neurons in the vlPBN, dlPBN, and elPBN. CGRP immunohistochemistry delineates the elPBN. Scale bars, 50 μm. (F-M) mCherry-labeled axons were detectable in the CeA, BNST, PVH, DMH, and ARC. Scale bars, 50 μm. (N) Strategy for selective labeling of DMH-projecting LPBN^BDNF^ neurons in *Bdnf^2A-Cre/+^* mice with mCherry. (O) EGFP marks the injection site in the DMH. Scale bar, 50 μm. (P-R) mCherry-labeled DMH-projecting LPBN^BDNF^ neurons in the vlPBN, dlPBN, and elPBN. CGRP immunohistochemistry delineates the elPBN. Scale bars, 50 μm. (S-W) mCherry-labeled axons from DMH-projecting LPBN^BDNF^ neurons were detected in the PVH (U), DMH (V), and ARC (W), whereas little or no axonal labeling was observed in the CeA (S) or BNST (T). Scale bars, 50 μm. Abbreviations: c, central lateral PBN; d, dorsal lateral PBN; e, external lateral PBN; v, ventral lateral PBN; m, medial PBN; scp, superior cerebellar peduncle; ARC, arcuate nucleus; BLA, basolateral amygdala; BNST, bed nucleus of the stria terminalis; CeA, central amygdala; DMH, dorsomedial hypothalamus; PVH, paraventricular hypothalamus.

Because a small subset of PVH-innervated LPBN^BDNF^ neurons was located in the elPBN and co-expressed CGRP (Figures 6C-6E), these neurons could account for the sparse projections to the CeA and BNST. Consistent with this possibility, selectively labeled PVH-innervated elPBN^CGRP^ neurons projected to both regions (Figure S7).

To selectively examine the major vl/dlPBN^BDNF^ population while minimizing inclusion of elPBN^CGRP^ neurons, we retrogradely labeled DMH-projecting LPBN^BDNF^ neurons by injecting AAV2retro-Ef1a-DIO-FLPo into the DMH and FLPo-dependent AAV2-Ef1a-fDIO-mCherry into the LPBN of *Bdnf^2A-Cre/+^*mice (Figures 6N-6R). Labeled neurons were concentrated in the vlPBN and dlPBN, with substantially fewer in the elPBN (Figure 6P). These neurons projected to the DMH, PVH, and ARC but showed little or no innervation of the CeA or BNST (Figures 6S-6W), indicating that DMH-projecting vl/dlPBN^BDNF^ neurons collateralize to multiple hypothalamic targets.

We obtained similar results by retrogradely labeling PVH-projecting LPBN^BDNF^ neurons (Figure S8). These neurons projected to the PVH, DMH and ARC but showed little or no innervation of the CeA or BNST (Figures S8F-S8J). Together, these complementary tracing experiments demonstrate that vl/dlPBN^BDNF^ neurons preferentially innervate a network of hypothalamic regions, whereas elPBN^CGRP^ neurons primarily target the CeA, BNST, VPMpc, and PSTN. These distinct projection patterns support functionally divergent roles for the two neuronal populations in the regulation of feeding.

### Activation of vl/dlPBN^BDNF^ neurons induces non-aversive feeding suppression

To examine the role of vl/dlPBN^BDNF^ neurons in feeding regulation, we bilaterally injected 8-week-old *Bdnf^2A-Cre/+^* mice with AAV2retro-Ef1a-DIO-FLPo and AAV2-CMV-EGFP into the DMH and either AAV5-Ef1a-fDIO-mCherry or AAV5-hSyn-fDIO-hM3D(Gq)-mCherry into the LPBN (Figure 7A). This strategy labeled the DMH injection site with EGFP (Figure 7B1) and enabled relatively selective expression of either mCherry or hM3D(Gq) predominantly in vl/dlPBN^BDNF^ neurons (Figures 7B2 and 7B3). CNO administration robustly suppressed nocturnal food intake in mice expressing hM3D(Gq), but not in control mice expressing mCherry (Figure 7C). To determine whether this feeding suppression was associated with aversion, we performed a saccharin conditioned taste aversion (CTA) assay in which consumption of a novel saccharin solution was paired with CNO-induced activation of vl/dlPBN^BDNF^ neurons. In a subsequent two-bottle preference test, hM3D(Gq)-expressing mice showed no reduction in saccharin preference compared with control mice (Figure 7D), indicating that activation of vl/dlPBN^BDNF^ neurons did not confer negative valence to the paired taste. Thus, unlike activation of aversive feeding circuits, chemogenetic activation of vl/dlPBN^BDNF^ neurons suppresses food intake without inducing CTA.

**Figure 7.**
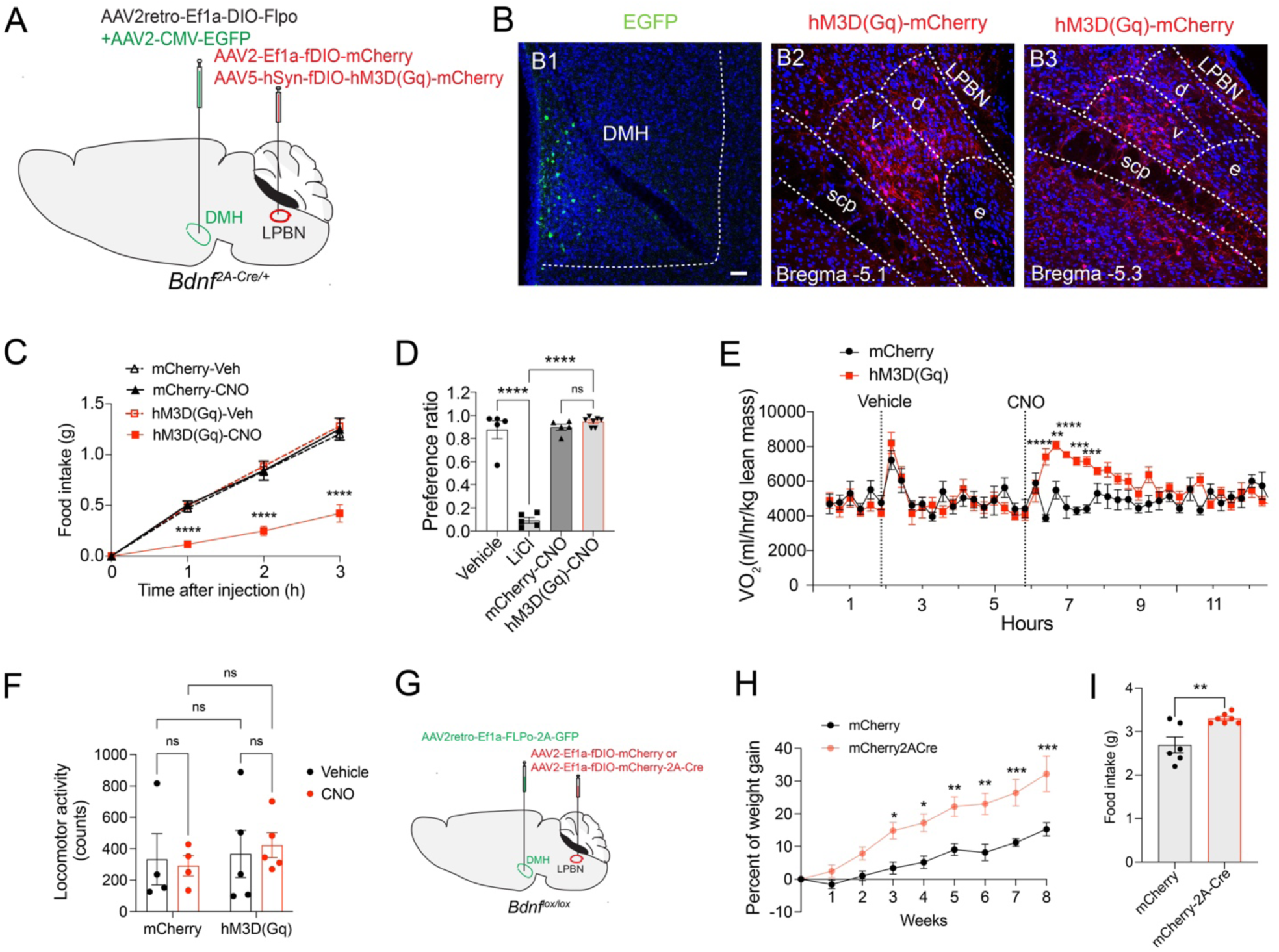
Activation of PVH-innervated vl/dlPBN^BDNF^ neurons induced non-aversive feeding suppression via a mechanism dependent on BDNF-TrkB signaling. (A) Strategy for selective expression of either mCherry or hM3D(Gq)-mCherry in vl/dlPBN^BDNF^ neurons. (B) EGFP labeled the injection site in the DMH (B1), whereas mCherry labeled mCherry-or hM3D(Gq)-expressing neurons in the LPBN at two rostrocaudal levels (B2 and B3). Scale bar, 50 μm. (C) CNO administration suppressed nocturnal food intake in mice expressing hM3D(Gq) in vl/dlPBN^BDNF^ neurons. n=6-9 mice per group. Two-way RM ANOVA followed by Bonferroni’s multiple comparisons test, ****p< 0.0001 when compared with vehicle treatment. (D) Activation of vl/dlPBN^BDNF^ neurons did not cause saccharin-conditioned taste aversion. n=5-7 mice per group. One-way ANOVA followed by Bonferroni’s multiple comparisons test, ns = not significant and ****p< 0.0001. (E) Activation of vl/dlPBN^BDNF^ neurons increased oxygen consumption (VO_2_). n=4-6 mice per group. Two-way ANOVA followed by Bonferroni’s multiple comparisons test, *p<0.05, **p<0.01, and ****p< 0.0001. (F) Average locomotor activity during the 2 h following treatment. n=4-5 mice per group. Two-way ANOVA followed by Fisher’s LSD post hoc test: ns, not significant. (G) Strategy for selective *Bdnf* deletion in DMH-projecting vl/dlPBN neurons. (H and I) Female mice with *Bdnf* deletion in the vl/dlPBN (mCherry-2A-Cre mice) exhibited increased body weight gain and hyperphagia, compared with control mice (mCherry mice). n=6 mice per group. Body weight: two-way ANOVA followed by Sidak’s multiple comparisons test, *p<0.05, **p<0.01, and ***p< 0.001; food intake: unpaired t test, **p<0.01. All data are presented as mean ± SEM.

In addition, activation of these neurons increased oxygen consumption without altering locomotor activity (Figures 7E and 7F). Because activation of LPBN-projecting PVH^TrkB^ neurons did not affect energy expenditure (Figure 2D), these findings suggest that vl/dlPBN^BDNF^ neurons integrate additional afferent inputs beyond PVH^TrkB^ neurons to regulate energy expenditure.

We next tested whether BDNF produced by these neurons is required for feeding regulation by deleting *Bdnf* selectively in DMH-projecting vl/dlPBN^BDNF^ neurons. To achieve this, AAV2retro-Ef1a-FLPo-2A-GFP was injected into the DMH and AAV2-Ef1a-mCherry-2A-Cre into the LPBN of 8-week-old *Bdnf^flox/flox^* mice (Figure 7G). Control mice received AAV2-Ef1a-fDIO-mCherry in the LPBN. Efficient *Bdnf* deletion was confirmed by mCherry immunohistochemistry and FISH for *Bdnf* mRNA (Figures S9A and S9B). Approximately 80% of DMH-projecting vlPBN neurons expressed *Bdnf* mRNA in control mice, whereas fewer than 5% of mCherry-labeled neurons expressed *Bdnf* in Cre-injected mice (Figure S9C), confirming efficient deletion. Loss of *Bdnf* in these neurons increased both food intake and body weight gain (Figures 7H and 7I). Together, these findings demonstrate that vl/dlPBN^BDNF^ neurons mediate non-aversive feeding suppression and that this function depends, at least in part, on BDNF-TrkB signaling.

### Presynaptic inputs to vl/dlPBN^BDNF^ neurons

To identify presynaptic inputs to vl/dlPBN^BDNF^ neurons, we performed monosynaptic retrograde tracing using a modified rabies virus system^50–54^. AAV2-CMV-FLEX-TVAmCherry-2A-oG^55,56^ was injected into the LPBN of *Bdnf^2A-Cre/+^* mice to express TVA-mCherry and optimized rabies glycoprotein (oG) selectively in LPBN^BDNF^ neurons (Figures 8A and 8B). Three weeks later, SADΔG-EGFP(EnvA) was injected into the DMH, where it infected the axon terminals of DMH-projecting vl/dlPBN^BDNF^ neurons and spread monosynaptically to their direct presynaptic partners^57,58^. Mice were perfused 10 days later, and EGFP-labeled presynaptic neurons were identified in coronal brain sections (Figure 8C).

**Figure 8.**
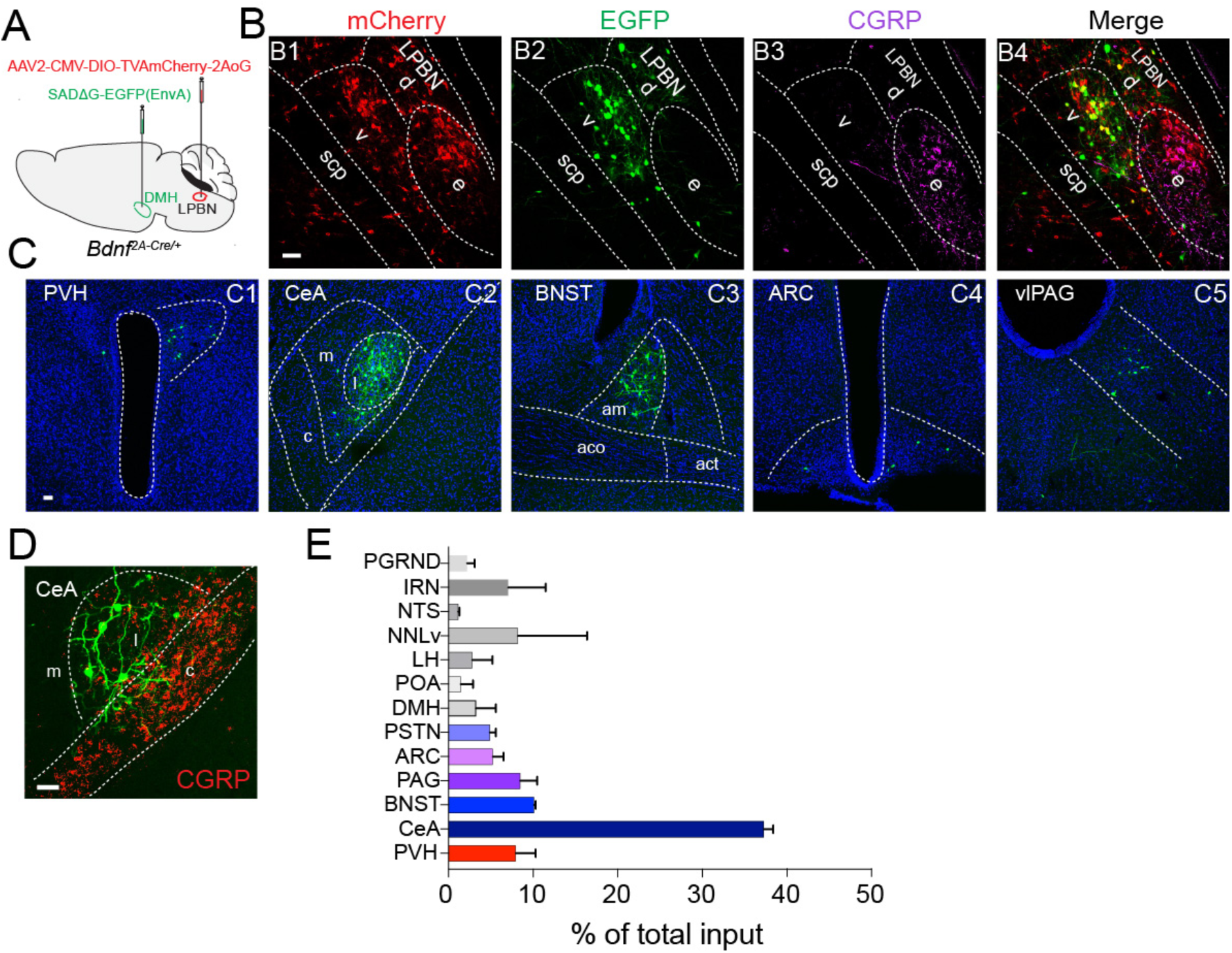
Presynaptic inputs to vl/dlPBN^BDNF^ neurons. (A) Virus injection into *Bdnf^2A-Cre/+^* mice. (B) Virus transduction in the LPBN. Scale bar, 50 μm. Abbreviations: v, vlPBN; e, elPBN. (C) EGFP-labeled presynaptic neurons in the PVH, CeA, BNST, ARC, and vlPAG. Scale bar, 50 μm. Abbreviations: l, lateral central amygdala; am, anteromedial area of the BNST. (D) A confocal image showing the locations of the presynaptic neurons and CGRP^+^ axonal terminals in the CeA. Scale bar, 50 μm. Abbreviations: c, capsular division of the CeA; l, lateral division of the CeA; m, medial division of the CeA. (E) Quantification of presynaptic neurons. N=3 mice. Abbreviations: PGRND, paragigantocellular reticular nucleus dorsal part; IRN, intermediate reticular nucleus; NNLv, nucleus of lateral lemniscus ventral part; LH, lateral hypothalamus; vlPAG, ventrolateral periaqueductal gray.

Quantitative analysis revealed that the CeL provides the largest presynaptic input to vl/dlPBN^BDNF^ neurons (Figures 8C2, 8D, and 8E). Additional major inputs arose from brain regions implicated in feeding regulation, including the PVH, ARC and DMH^2^. Inputs from thermoregulatory regions, such as the DMH and preoptic area (POA)^61^, may contribute to the increase in energy expenditure elicited by activation of vl/dlPBN^BDNF^ neurons (Figure 7E). Notably, these neurons receive very little direct input from the NTS (Figure 8D). Instead, many of their major presynaptic input regions, including the CeL, PSTN, and BNST, are known postsynaptic targets of elPBN^CGRP^ neurons (Figures 8D and S6). These findings suggest that noxious stimuli may influence vl/dlPBN^BDNF^ neurons indirectly through elPBN^CGRP^ neurons and their downstream targets, thereby integrating aversive and homeostatic signals to regulate feeding.

## DISCUSSION

This study demonstrates that PVH^TrkB^ neurons project to BDNF-expressing neurons located predominantly in the vlPBN and dlPBN to suppress feeding without eliciting aversion. BDNF-expressing neurons constitute the largest molecularly defined neuronal population in these PBN subdivisions and were preferentially recruited by refeeding-associated satiety signals. Although BDNF expression partially overlaps with Foxp2 and Pdyn in the LPBN, chemogenetic activation of PVH-innervated LPBN^BDNF^ neurons, but not LPBN^Foxp2^ or LPBN^Pdyn^ neurons, suppressed feeding. This anorexigenic effect depended on BDNF-TrkB signaling and was required for long-term regulation of energy homeostasis. In addition, vl/dlPBN^BDNF^ neurons project to multiple hypothalamic nuclei, providing an anatomical substrate for coordinating feeding and energy expenditure. Together, these findings identify vl/dlPBN^BDNF^ neurons as a previously unrecognized PBN population that mediates non-aversive feeding suppression.

Nearly all LPBN-projecting, satiety-promoting PVH^MC4R^ neurons express TrkB, suggesting that vl/dlPBN^BDNF^ neurons are key downstream mediators of the anorexigenic effects of melanocortin signaling. Interestingly, vl/dlPBN^BDNF^ neurons receive prominent inputs from several brain regions, including the CeA, BNST, and PSTN, that are major postsynaptic targets of aversive elPBN^CGRP^ neurons, yet they are only weakly activated by LPS. This anatomical organization suggests that noxious stimuli may indirectly modulate vl/dlPBN^BDNF^ neuronal activity through the elPBN^CGRP^ circuit and its downstream targets, thereby allowing aversive signals to modestly influence homeostatic feeding circuits.

### vl/dlPBN^BDNF^ neurons mediate non-aversive feeding suppression

BDNF expression in the LPBN is concentrated in the elPBN, vlPBN, and dlPBN subdivisions. Because elPBN^CGRP^ neurons mediate aversive feeding suppression^4^ and BDNF expression largely overlaps with CGRP expression in the elPBN, elPBN^BDNF^ neurons are also likely to participate in aversive feeding suppression. In contrast, although the vlPBN and dlPBN are anatomically adjacent to the elPBN and express lower levels of BDNF, our findings provide multiple lines of evidence that vl/dlPBN^BDNF^ neurons constitute a functionally distinct population that mediates non-aversive feeding suppression.

First, vl/dlPBN^BDNF^ neurons receive direct input from PVH^TrkB^ and PVH^MC4R^ neurons. TrkB and MC4R are essential regulators of energy homeostasis, and loss-of-function mutations in either receptor cause hyperphagia and severe obesity in humans and mice^33,62^. Moreover, the PVH-to-LPBN pathway mediates part of the anorexigenic action of these signaling systems, and optogenetic activation of LPBN-projecting PVH^MC4R^ neurons induces real-time place preference^24,26^. These observations indicate that vl/dlPBN^BDNF^ neurons are embedded within hypothalamic circuits that regulate homeostatic feeding rather than aversive responses.

Second, refeeding selectively increased *Bdnf* expression in the vlPBN and induced *Fos* more strongly in the vlPBN than in the elPBN, whereas the aversive stimulus LPS preferentially activated the elPBN, consistent with previous findings^5^. These distinct activation patterns suggest that vl/dlPBN^BDNF^ neurons preferentially respond to physiological satiety signals, whereas elPBN^CGRP/BDNF^ neurons are preferentially recruited by noxious stimuli. This functional distinction is further supported by their afferent connectivity. Whereas elPBN^CGRP^ neurons receive dense input from the NTS, a major relay for visceral and noxious sensory information^4^, our monosynaptic tracing revealed that vl/dlPBN^BDNF^ neurons receive little direct input from the NTS.

Third, chemogenetic activation of vl/dlPBN^BDNF^ neurons suppressed feeding without inducing CTA or altering locomotor activity, indicating that these neurons suppress feeding without eliciting defensive or anxiety-related behaviors typically associated with aversive stimuli^63^.

Finally, the PVH^TrkB^-to-vl/dlPBN^BDNF^ circuit was both sufficient and necessary for feeding suppression, whereas silencing aversive elPBN^CGRP^ neurons does not increase food intake^7^. Together, these findings identify vl/dlPBN^BDNF^ neurons as the LPBN population that mediates physiological, non-aversive feeding suppression.

Although PVH neurons predominantly innervate vl/dlPBN^BDNF^ neurons, they also project to a small subset of elPBN^CGRP/BDNF^ neurons. Given the established role of elPBN^CGRP^ neurons in mediating aversive feeding suppression, activation of this minor PVH-to-elPBN pathway may also contribute to aversive suppression of feeding. However, activation of all PVH-innervated LPBN^BDNF^ neurons failed to induce CTA, suggesting that activation of this pathway alone is insufficient to elicit aversion or that any aversive effect is outweighed by the positive valence generated by concurrent activation of vl/dlPBN^BDNF^ neurons. It will be interesting to determine whether this PVH-to-elPBN pathway contributes to the unpleasant physiological or psychological consequences of excessive food consumption, thereby providing a negative-feedback mechanism that limits overeating.

### BDNF is a marker for appetite-controlling neurons in the vlPBN and dlPBN

Our FISH analysis revealed that refeeding following an overnight fast robustly induced *Fos* expression in the vlPBN and, to a lesser extent, in the dlPBN. In the vlPBN, 88% of *Fos*-positive cells expressed *Bdnf*, whereas 73% of *Bdnf*-expressing neurons were positive for *Fos*. These findings indicate that *Bdnf* is a useful molecular marker for LPBN neurons involved in non-aversive feeding suppression. However, it is not an exclusive marker, because not all refeeding-activated neurons express *Bdnf* and *Bdnf* is also expressed in other LPBN populations, including aversion-associated elPBN^CGRP^ neurons.

*Foxp2* and *Pdyn* identify two additional LPBN neuronal populations implicated in feeding regulation^59,60^. However, chemogenetic activation of PVH-innervated LPBN^Foxp2^ or LPBN^Pdyn^ neurons failed to suppress nocturnal food intake, despite the observation that refeeding activated a large proportion of both populations, as indicated by *Fos* induction. These findings suggest that activation of either molecularly defined population alone is insufficient to drive feeding suppression.

One explanation is that the neurons mediating non-aversive feeding suppression are distributed across multiple molecularly defined populations, with *Bdnf* labeling a substantially larger fraction than either *Foxp2* or *Pdyn*. Alternatively, the critical satiety-promoting neurons may be defined by additional molecular markers that remain to be identified. Consistent with this possibility, recent single-cell transcriptomic analyses of the PBN did not identify a neuronal cluster selectively localized to the vlPBN and dlPBN^64–67^, suggesting that greater transcriptomic depth and/or spatial resolution may be required to resolve this functionally distinct population. Future single-cell transcriptomic analyses of the PBN in fasted and refed mice, ideally coupled with activity-dependent labeling and functional circuit interrogation, should facilitate the identification of more selective molecular markers for this neuronal population, thereby enabling more precise mechanistic studies of the neural circuits underlying non-aversive feeding suppression.

### Noxious stimuli engage vl/dlPBN^BDNF^ neurons through indirect pathways

Noxious stimuli activate elPBN^CGRP^ neurons through spinal and vagal afferent pathways^4,5,8,12^. For example, visceral malaise induced by food poisoning activates specific neuronal populations in the NTS via vagal afferents, which in turn provide direct excitatory input to elPBN^CGRP^ neurons^13^. Interestingly, our monosynaptic rabies tracing revealed that vl/dlPBN^BDNF^ neurons receive little direct input from the NTS but strong presynaptic input from regions innervated by elPBN^CGRP^ neurons, including the CeA, BNST, and PSTN. Nevertheless, intraperitoneal LPS administration, which mimics bacterial infection and produces sickness behavior including anorexia, weakly activated vl/dlPBN^BDNF^ neurons, as indicated by *Fos* induction. These observations suggest that noxious signals may reach vl/dlPBN^BDNF^ neurons indirectly through downstream targets of elPBN^CGRP^ neurons. The organization of the CeA provides a potential mechanism for such communication.

We found that elPBN^CGRP^ neurons densely innervated the CeC densely but much more sparsely innervated the CeL, whereas CeA neurons presynaptic to vl/dlPBN^BDNF^ neurons were localized almost exclusively in the CeL. Two circuit mechanisms could explain how this organization influences vl/dlPBN^BDNF^ neurons. First, PKCδ-expressing GABAergic neurons in the CeC and CeL receive monosynaptic input from elPBN^CGRP^ neurons^68^. If these neurons directly innervate vl/dlPBN^BDNF^ neurons, activation of the elPBN^CGRP^ pathway by noxious stimuli would inhibit rather than activate vl/dlPBN^BDNF^ neurons. Such inhibition could limit recruitment of this homeostatic feeding-suppression circuit during illness or threat.

Alternatively, because CeL neurons are GABAergic^69^, the weak activation of vl/dlPBN^BDNF^ neurons by LPS could arise through disinhibition. One possible pathway is: LPS ◊ NTS ◊ elPBN^CGRP^ ◊ CeL PKCδ^+^ ⊣ CeL neurons ⊣ vl/dlPBN^BDNF^ neurons. In this model, activation of PKCδ^+^ neurons inhibits a second population of GABAergic CeL neurons that normally suppress vl/dlPBN^BDNF^ neurons, resulting in disinhibition of vl/dlPBN^BDNF^ neurons. Although this circuit remains hypothetical, it is consistent with our anatomical tracing data and the known organization of CeA microcircuits. Moreover, the polysynaptic nature of this pathway, together with the relatively sparse innervation of the CeL by elPBN^CGRP^ neurons, could explain why LPS activates vl/dlPBN^BDNF^ neurons much less strongly than physiological refeeding.

Recent single-cell RNA-seq studies indicate that the CeL and CeC contain more than ten transcriptionally distinct inhibitory neuronal populations^70,71^. Future studies combining projection-specific monosynaptic rabies tracing with multiplex FISH or spatial transcriptomics should identify the molecular identity of CeL neurons that project to vl/dlPBN^BDNF^ neurons. Functional manipulation of these defined CeL populations will determine whether CeA circuits inhibit or disinhibit vl/dlPBN^BDNF^ neurons during aversive states and reveal how aversive signals interact with homeostatic feeding circuits to regulate food intake.

### How does BDNF-TrkB signaling modulate the PVH-to-LPBN pathway?

Our findings indicate that neuronal activity and BDNF-TrkB signaling reinforce one another within the anorexigenic PVH-to-LPBN circuit. Whole-cell patch-clamp recordings showed that refeeding increased the frequency of spontaneous EPSCs in LPBN-projecting PVH^TrkB^ neurons, indicating enhanced excitatory synaptic input that is expected to increase neuronal firing. Consistent with this interpretation, refeeding also induced *Fos* expression in these neurons^26^. Increased neuronal activity is known to stimulate *Bdnf* transcription and promote BDNF secretion^47^. Accordingly, we found that refeeding increased *Bdnf* mRNA levels in vl/dlPBN neurons. In cortical and hippocampal circuits, BDNF enhances glutamatergic synaptic transmission and promotes synaptic potentiation^72–76^. Therefore, the refeeding-induced increase in mEPSC amplitude in LPBN-projecting PVH^TrkB^ neurons is likely mediated, at least in part, by enhanced BDNF-TrkB signaling. Such activity-dependent synaptic potentiation may establish a positive-feedback mechanism that amplifies the anorexigenic output of the PVH-to-LPBN circuit during satiety. Consistent with this model, the selective TrkB inhibitor ANA-12 significantly attenuated the anorexigenic effect produced by chemogenetic activation of either LPBN-projecting PVH^TrkB^ neurons or vl/dlPBN^BDNF^ neurons.

Nearly all LPBN-projecting PVH neurons express TrkB, whereas the vast majority of PVH-innervated neurons in the vl/dlPBN express BDNF. Gene deletion experiments further demonstrate the importance of this BDNF-TrkB signaling pathway in energy homeostasis, as *Ntrk2* deletion in LPBN-projecting PVH neurons and *Bdnf* deletion in vl/dlPBN neurons resulted in hyperphagia and obesity in our previous and current studies, repectively^26^. Because BDNF is produced postsynaptically in vl/dlPBN neurons, these findings raise the possibility that target-derived BDNF is released from vl/dlPBN neurons and acts retrogradely on TrkB receptors expressed by LPBN-projecting PVH neurons to strengthen excitatory synaptic transmission. This model is supported by our observation that refeeding increased the amplitude of mEPSCs in LPBN-projecting PVH^TrkB^ neurons, an effect that is consistent with enhanced BDNF-TrkB signaling. Although long-distance retrograde neurotrophic signaling is well established in the peripheral nervous system, where it is essential for neuronal survival^77^, BDNF signaling at central synapses is generally thought to occur through local autocrine/paracrine mechanisms or through anterograde release from presynaptic neurons^78,79^. If confirmed, our findings would reveal a previously unrecognized mode of BDNF signaling in which target-derived BDNF retrogradely modulates synaptic function at long-range hypothalamic projections.

## MATERIALS AND METHODS

### Animals

*Ntrk2^2A-Cre/+^ and* floxed *Bdnf* (*Bdnf^lox^*) mouse strains were previously described. *Bdnf^lox^* mice were backcrossed to a C57BL/6J background for at least 5 generations prior to use in this study. *Mc4r^2A-Cre/+^* (stock No: 030759)^24^*, GFP-L10a/+* (stock No: 024750)^44^, *Bdnf^2A-Cre/+^*(stock No: 030189)^80^, *Foxp2^Cre/+^*(stock No: 030541)^81^, *Pdyn^IRES-Cre/+^*(stock No: 27958)^82^, *Calca^Cre/+^*(stock No: 033168)^5^, and C57BL6/J (stock No: 000664) mouse strains were obtained from the Jackson Laboratory. *Mc4r^2A-Cre/+^;GFP-L10/+* mice were generated by crossing *Mc4r^2A-Cre/+^* mice with *GFP-L10a/+* reporter mice. All mice used in this study were maintained in the C56BL6/J genetic background. Mice were maintained on a 12-h/12-h light/dark cycle with *ad libitum* access to water and regular laboratory chow (Teklad Rodent Diet 2019; 3.3 kcal/g energy density, 22% kcal from fat) unless otherwise specified. Animals of both sexes were included in each experimental groups, and the data were combined for analysis. All experiments were performed in accordance with relevant guidelines and regulations regarding the use of experimental animals. All protocols were approved by the institutional Animal Care and Use Committees at the Herbert Wertheim UF Scripps Institute for Biomedical Innovation & Technology, University of Florida.

### Viruses

Plasmid pAAV-EF1a-DIO-FLPo-WPRE-hGHpA (#87306; a gift of Li Zhang)^49^ and pAAV-CMV-FLEX-TVAmCherry-2A-oG (#102985; a gift of Marco Tripodi)^55^ were obtained from Addgene. AAV2retro-DIO-FLPo (2.6 x 10^13^ vg/ml) and AAV2-CMV-FLEX-TVAmCherry-2A-oG (5.23 x 10^12^ vg/ml) virus were packaged by ViGene Biosciences Inc. AAV2retro-Ef1a-DIO-mCherry (4 x 10^12^ vg/ml), AAV2retro-hSyn-mCherry (5.3 x 10^12^ vg/ml), AAV5-hSyn-fDIO-hM3D(Gq)-mCherry (4.1 x 10^12^ vg/ml), AAV5-hSyn-fDIO-hM4D(Gi)-mCherry (4 x 10^12^ vg/ml), AAV2-CAG-FLEX-tdTomato (4.4 x 10^12^ vg/ml), and AAV2-CMV-GFP (4.4 x 10^12^ vg/ml) were purchased from UNC Vector Core. AAV1-Ef1a-DIO-Flpo (7 x 10^12^ vg/ml) and AAV8-hSyn-DIO-hM4Di-mCherry (2.3 x 10^13^ vg/ml) were purchased from Addgene. AAV2-Ef1a-fDIO-mCherry (4.2 x 10^12^ vg/ml), AAV2-Ef1a-fDIO-mCherry-P2A-Cre (2.1 x 10^12^ vg/ml), AAV2retro-hSyn-FLPo-T2A-GFP (6.9 x 10^12^ vg/ml), and RV-SADΔG-GFP(EnvA) (1.5 x 10^8^ IU/mL) were described previously^26,83^.

### Chemicals

The following chemicals were used in this study: clozapine N-oxide (CNO; Cayman Chemical, #25780), deschloroclozapine (DCZ; Tocris Bioscience, #7193,), ANA-12 (MedChemExpress, #HY-12497), Cremophor EL (EMD Millipore, #23847), Saccharin (Thermo Scientific, #149001000), and lithium chloride (Sigma Aldrich, #310468).

### Fluorescent in situ hybridization

Freshly dissected mouse brains were rapidly frozen in a 2-methylbutane (Fisher Scientific, #O3551-4) dry ice bath. Cryostat sections (14 µm thick) were collected on Superfrost Plus slides (Fisherbrand, #12-550-15) and processed for fluorescent in situ hybridization using the RNAscope Multiplex Fluorescent Reagent Kit v2 (Advanced Cell Diagnostics, #323100). Brain sections were fixed with pre-chilled 10% formalin at 4°C for 15 min and dehydrated through a graded ethanol series (50%, 75% and 100%). Sections were air-dried for 20 min, pretreated with Protease IV for 30 min, and washed twice in PBS. Sections were then incubated with the target probes for 2 hours at 40°C. The following probes were used: *mCherry* (#431201-C3), *Gfp* (#400281-C2), *Bdnf* (#424821-C4), *Ntrk2* (#423611-C3), *Slc17a6* (#319171-C3), *Slc32a1* (#319191-C2), *Foxp2* (#428791-C2), *Pdyn* (#318771-C3), *Fos* (#316921-C1), *Glp1r* (#418851), and *Calca* (#420361-C2). Signals were amplified by sequential incubation with v2Amp1 (30 min), v2Amp2 (30 min), and v2Amp3 (15 min), followed by incubation with TSA Plus Fluorescein/Cyanine5 (Akoya Biosciences, #NEL754001KT) or ClariTSA Fluorophore Kit 570 (Bio-Techne, #7526/1) for 30 min at 40°C. Slides were counterstained with DAPI, and images were acquired using a Nikon C2+ confocal microscope.

### Dual immunohistochemistry and fluorescent in situ hybridization

Mice were transcardially perfused with ice-cold phosphate buffered saline (PBS; Sigma-Aldrich, #P2287) and then 4% paraformaldehyde (PFA), and brains were post-fixed for 24 hours at 4°C. Brains were cryoprotected sequentially in 10%, 20%, and 30% sucrose for 24 h each at 4°C, followed by rapid freezing in a 2-methylbutane/dry ice bath. Frozen brains were embedded in Tissue-Tek Optimal Cutting Temperature (OCT) compound (Sakura Finetek, #4583), and 14-µm cryostat sections were collected onto Superfrost Plus slides for dual immunohistochemistry and fluorescent in situ hybridization (FISH). FISH was performed using the RNAscope Multiplex Fluorescent Reagent Kit v2 (Advanced Cell Diagnostics, #323100). Sections were fixed with pre-chilled 10% formalin at 4°C for 15 min and then dehydrated through a graded ethanol series (50%, 70%, and 100%). After air-drying for 20 min, sections were treated with hydrogen peroxide for 10 min, rinsed twice in distilled water, and incubated in target retrieval buffer at 95°C for 5 min. Sections were rinsed twice in distilled water and briefly washed in 1× PBST (0.005% Tween-20 in PBS). Sections were incubated overnight at 4°C with a Living Colors DsRed polyclonal antibody (Takara Bio, #632496; 1:500) or chicken anti-GFP antibody (Abcam, #ab13970; 1:1000) in Codetection Antibody Diluent (Advanced Cell Diagnostics, #323160). On following day, sections were washed three times in 1× PBST, post-fixed with 10% formalin for 30 min at room temperature, and washed four additional times in 1× PBST. Protease Plus was then applied for 30 min at 40°C, followed by two PBS washes. Sections were hybridized with either Mm-Bdnf-C4 (Advanced Cell Diagnostics, #424821-C4) or Mm-Ntrk2-C3 (Advanced Cell Diagnostics, #423611-C3) for 2 h at 40°C. Signal amplification was performed by sequential incubation with v2Amp1 (30 min), v2Amp2 (30 min), v2Amp3 (15 min), and v2C4-HRP (15 min), followed by TSA Plus Fluorescein (Akoya Biosciences, #NEL741001KT,) or ClariTSA Fluorophore Kit 570 (Bio-Techne, #7526/1) for 30 min at 40°C. After three washes in 1× PBST, sections were incubated for 1 h at room temperature with anti-rabbit Alexa Fluor 594 or anti-chicken Alexa Fluor 488 secondary antibody (Jackson ImmunoResearch, 1:800). Slides were counterstained with DAPI, and images were acquired using a Nikon C2+ confocal microscope.

### Sample preparation for the Fos measurement

For fasting and refeeding experiments, four mice (2 male and 2 female) were used per group. One week before sample collection, mice were transferred to a thermoneutral room (30°C) and handled daily in their home cages to minimize temperature- and handling-induced stress responses. All animals were food-deprived overnight (15 h). Mice in the fasted group were euthanized immediately after the fast, whereas those in the refed group were given access to chow pellets for 2 h and were euthanized immediately afterward.

For LPS treatment, four mice (2 male and 2 female) were used per group. Mice were housed at room temperature and handled daily in their home cages for one week before the experiment. After the handling period, mice were food-deprived overnight. The following day, they were randomly assigned to groups and injected intraperitoneally with vehicle and LPS (100 µg/kg). Mice were euthanized 90 min after injection, and brains were collected for in situ hybridization.

### Quantification of *Fos* and *Bdnf* expression

For FISH, brains were sectioned coronally at a thickness of 14 µm. Ten series of PVH-containing sections and seven series of LPBN-containing sections were collected, with each series consisting of one slide containing 4–6 sections. One slide from each region was used for each FISH experiment. The analyzed sections spanned the PVH from Bregma −0.58 mm to −1.06 mm and the LPBN from Bregma −5.0 mm to −5.4 mm. All analyses were conducted using NIS-Elements AR software (Nikon). Co-expression of mRNA signals was quantified in DAPI-positive cells within the PVH and LPBN. Cells containing two or more puncta for a given transcript were considered positive. To quantify *Bdnf* and *Fos* signals, a region of interest (ROI) was drawn around the vlPBN and elPBN at matched rostrocaudal levels, and fluorescence intensity for each transcript was measured. Background fluorescence measured in a cell-free region within the same section, was subtracted from each measurement.

### Immunohistochemistry

Mice were deeply anaesthetized with avertin and transcardially perfused with PBS, followed by 4% paraformaldehyde in PBS. Coronal brain sections (40 μm in thickness) were obtained using a sliding microtome (Leica SM2000R). Brain sections were rinsed once with Tris-buffered saline (TBS; 10 mM Tris-HCl, 150 mM NaCl, pH 7.5), incubated with blocking buffer (0.4% Triton X-100, 1% bovine serum albumin, and 10% goat serum or horse serum in TBS) for 1 h at room temperature, and then incubated with primary antibodies diluted in the blocking buffer overnight at room temperature. The following primary antibodies were used: rabbit anti-DsRed (1:1000; TaKaRa, #632496), and goat anti-CGRP (1:1,000; Abcam). After three washes in TBS, sections were incubated with appropriate fluorescent secondary antibodies (1;500; Jackson ImmunoResearch) for 1 h at room temperature. Sections were washed three times in TBS, mounted onto slides, and coverslipped with a mounting medium containing 4′,6-diamidino-2-phenylindole (DAPI; Vector Laboratories). Fluorescent images were captured using a Nikon C2+ confocal microscope.

### Stereotaxic surgery

Mice (7-8 weeks old) were deeply anesthetized with isoflurane and placed onto a stereotaxic apparatus (David Kopf, Model 940). A small incision was made to expose the skull, and a small hole was drilled on skull above the injection site. A Nanofil 33-gauge needle (World Precision Instruments, #NF33BV-2) was inserted into the brain, and the virus was injected at a rate of 25 nl per min using a Microsyringe Pump (World Precision Instruments, Micro4). After infusion, the needle was kept in the same position for 5 min. Following injections, mice received a subcutaneous injection of Loxicom (1.5 mg per kg) to relieve pain. Stereotaxic coordinates from the bregma and skull used for this study were: PVH (AP: - 0.45 mm, ML: ±0.25 mm, DV: -5.4 mm), DMH (AP: -1.25 mm, ML: ±0.35 mm, DV: -5.64 mm), LPBN (AP: -5 mm, ML: ±1.5 mm, DV: -3.8 mm). After physiological measurements were completed, animals were perfused with PBS and then 4% paraformaldehyde, and their brains were sliced for examination of injection sites. Mice with missed injections were either assigned to a separate group or excluded from the final data analysis.

### Electrophysiology

Age-matched *Ntrk2^2A-Cre/+^* mice injected with AAV2retro-DIO-mCherry into the LPBN were used for whole-cell patch-clamp recordings and matched to cohorts used for feeding behavioral assays. Mice were deeply anesthetized with isoflurane, quickly decapitated, and the brains were dissected and placed in ice-cold artificial cerebrospinal fluid (aCSF) containing (in mM): 124 NaCl, 3 KCl, 26 NaHCO₃, 1.25 NaH₂PO₄, 1 MgSO₄, 2 CaCl₂, and 10 D-glucose, equilibrated with 95% O₂ and 5% CO₂. Coronal PVH slices (250 µm) were cut using a vibratome (Leica VT1200S, Germany) and transferred to oxygenated aCSF at 32°C for 30 min recovery, then maintained at room temperature (23–25°C) for an additional 30 min before recording^84,85^.

Slices were gently transferred to a recording chamber (RC-27, Warner Instruments, Hamden, CT) continuously perfused with oxygenated aCSF at 2–3 ml/min. LPBN-projecting TrkB neurons in the PVH were visually identified using an Olympus BX51WI microscope under brightfield and red fluorescence channels. Whole-cell voltage-clamp recordings were performed using borosilicate glass pipettes (World Precision Instruments, Sarasota, FL) with a resistance of 3–5 MΩ, pulled with a micropipette puller (P-1000, Sutter Instrument, Novato, CA).

For excitatory current recordings, pipettes were filled with internal solution containing (in mM): 115 CsMeSO₃, 20 CsCl, 10 HEPES, 0.6 EGTA, 4 Mg-ATP, 0.3 Na₃-GTP, 1 QX-314, 2.5 MgCl₂, and 10 Na₂-phosphocreatine (pH 7.3 with CsOH; osmolarity 285 mOsm). For inhibitory current recordings, the internal solution contained (in mM): 100 CsCl, 35 CsMeSO₃, 10 HEPES, 0.5 EGTA, 4 Mg-ATP, 0.3 Na₃-GTP, 5 QX-314, and 10 Na₂-phosphocreatine (pH 7.3 with CsOH; osmolarity 285 mOsm).

Miniature excitatory postsynaptic currents (mEPSCs), spontaneous excitatory postsynaptic currents (sEPSCs), and spontaneous inhibitory postsynaptic currents (sIPSCs) were recorded by holding neurons at −70 mV without series resistance or liquid junction potential compensation. Neurons with access resistance (Ra) > 25 MΩ were excluded. To isolate the currents, mEPSCs were recorded in the presence of 1 µM tetrodotoxin (TTX; American Radiolabeled Chemicals, # ARC D0640) and 100 µM picrotoxin (PTX; Sigma-Aldrich, # P1675); sEPSCs were recorded in the presence of 100 µM PTX; and sIPSCs were recorded in the presence of 20 µM CNQX (Tocris, # 1045) and 50 µM D-APV (Tocris, # 0106).

Signals were acquired using a Multiclamp 700B amplifier and DigiData 1550A digitizer (Molecular Devices, San Jose, CA). Data were low-pass filtered at 2.9 kHz, and sampled at 10 kHz. Analyses of mEPSCs, sEPSCs, and sIPSCs were performed using Clampfit 10.6. Synaptic events were detected using the template search method with a template matching threshold of 2, followed by manual verification of detected events. For each neuron, 100 events with amplitudes greater than 5 pA were used to calculate the average current amplitude and frequency.

### Physiological measurements

In the *Bdnf* gene deletion experiment, body weight was measured weekly. For food intake, mice were individually housed, and daily food intake was measured over 5 days after 3 days of acclimation. Body composition was determined using a Minispec LF-50/mq 7.5 NMR Analyzer (Brucker Optics). A comprehensive laboratory animal monitoring system (CLAMS; Columbus Instruments) was used to measure oxygen consumption (VO₂), carbon dioxide production (VCO₂), and locomotor activity. Mice were acclimated to the metabolic chambers for at least 2 days before experiments. To assess the effects of chemogenetic neuronal activation on energy expenditure, mice received an intraperitoneal injection of saline at 9:00 AM, followed by an injection of CNO (1 mg/kg; Cayman Chemical) at 1:00 PM.

### Chemogenetic manipulation of food intake

Mice expressing mCherry, hM3D(Gq)-mCherry, or hM4D(Gi)-mCherry were singly housed for at least one week with daily handling before the experiment. For food intake measurement during the light cycle, mice were transferred to a clean cage without food between 7:00-8:00 AM and injected intraperitoneally with vehicle or DCZ (100 µg/kg) one hour later. Pre-weighed food was introduced 30 minutes after injection, and food intake was measured hourly for 3 h. For food intake measurement during the dark cycle, mice were transferred to a clean cage without food three hours before lights off and injected with vehicle or CNO (1 mg/kg) 30 minutes before lights off. Pre-weighed food was provided at lights off, and food intake was measured hourly for the first three hours of the dark cycle. To block TrkB signaling, mice received an additional injection of ANA-12 (1.5 mg/kg in 2% DMSO and 20% Cremophor EL in saline) 30 min prior to CNO injection; all subsequent procedures were as described above.

### Rectal temperature measurements

Core body temperature was measured 1 hour after saline or CNO (1 mg/kg) injection, between 1:00 PM and 2:00 PM. A digital thermometer (Fisher Traceable Type K) with a rodent rectal probe (World Precision Instruments, RET-3) was used to obtain rectal temperature readings.

### Saccharin conditioned taste aversion

Mice were singly housed for at least 1 week with daily handling before the experiment. Water was delivered via two drinko measurers (leak-free water bottles; Amuza Inc.), and daily water intake was monitored to confirm stable acclimation to the apparatus. On Day 1 (conditioning session), water tubes were removed three hours before lights off. At lights off, two tubes containing 0.15% saccharin solution were provided for one hour, after which mice received an injection of LiCl (126 mg/kg), saline, or CNO (1mg/kg). The saccharin solution tubes were immediately replaced with regular water tubes following injection. On Day 3 (testing session), water tubes were again removed three hours before lights off. At lights off, mice were simultaneously presented with one tube containing 0.15% saccharin solution and one tube containing water, with tube positions randomized. Intake from both tubes was measured two hours after presentation. The preference ratio was calculated as: Preference Ratio = Saccharin solution intake / (Saccharin solution intake + Water intake).

### Monosynaptic retrograde tracing using modified rabies virus

AAV2-CMV-DIO-TVA-mCherry-2A-oG virus was unilaterally injected into the LPBN of *Bdnf2A-Cre*/+ mice. Three weeks later, SADΔG-GFP rabies virus was unilaterally injected into the ipsilateral DMH. Mice were allowed 7 days for retrograde transport of the rabies virus and GFP expression, then transcardially perfused with 4% paraformaldehyde in PBS as described above. Coronal whole-brain sections (50 µm) were collected on a sliding microtome (Leica SM2000R). Every other section from two injected mice was used for presynaptic neuron quantification. The percentage of inputs from each brain region was calculated by dividing the number of GFP-positive neurons in each brain region by the total number of GFP-positive neurons across the entire brain.

### Statistical analysis

Statistical analyses were performed using Prism 10 (GraphPad Software). All data are expressed as mean ± SEM. Data were tested using two-tailed unpaired or paired *t*-test or two-way ANOVA with post hoc Bonferroni’s or Tukey’s multiple comparisons for statistical significance. Experiments in this study were repeated independently two to three times. No statistical methods were used to determine the sample size for each experiment. The animal numbers were estimated based on our previous studies. Half of the brain sections from each virus-injected mouse were used for histological examination of injection sites. Mice with weak or off-target viral expression were excluded from final data analysis. Each micrograph shows a representative situation for each group of mice.

## ACKNOWLEDGMENTS

We thank Dr. Melissa Kazantzis for assistance in measurements of physical activity and oxygen consumption using CLAMS. This work was supported by grants the National Institutes of Health, United States to B.X. (R01 DK105954 and R01 DK134650).

## AUTHOR CONTRIBUTIONS

JJA and BX designed research; JJA, JWT, YC, and XY performed experiments; JJA and JWT analyzed data; and JJA and BX wrote the paper.

## Declaration of generative AI and AI-assisted technologies in the manuscript preparation process

During the preparation of this manuscript, the authors used ChatGPT to improve the clarity and readability of the text. The authors reviewed and edited the AI-assisted output as needed and take full responsibility for the content of the manuscript.

