## Supplemental figures for "A PVH-to-parabrachial BDNF circuit mediates non-aversive feeding suppression"

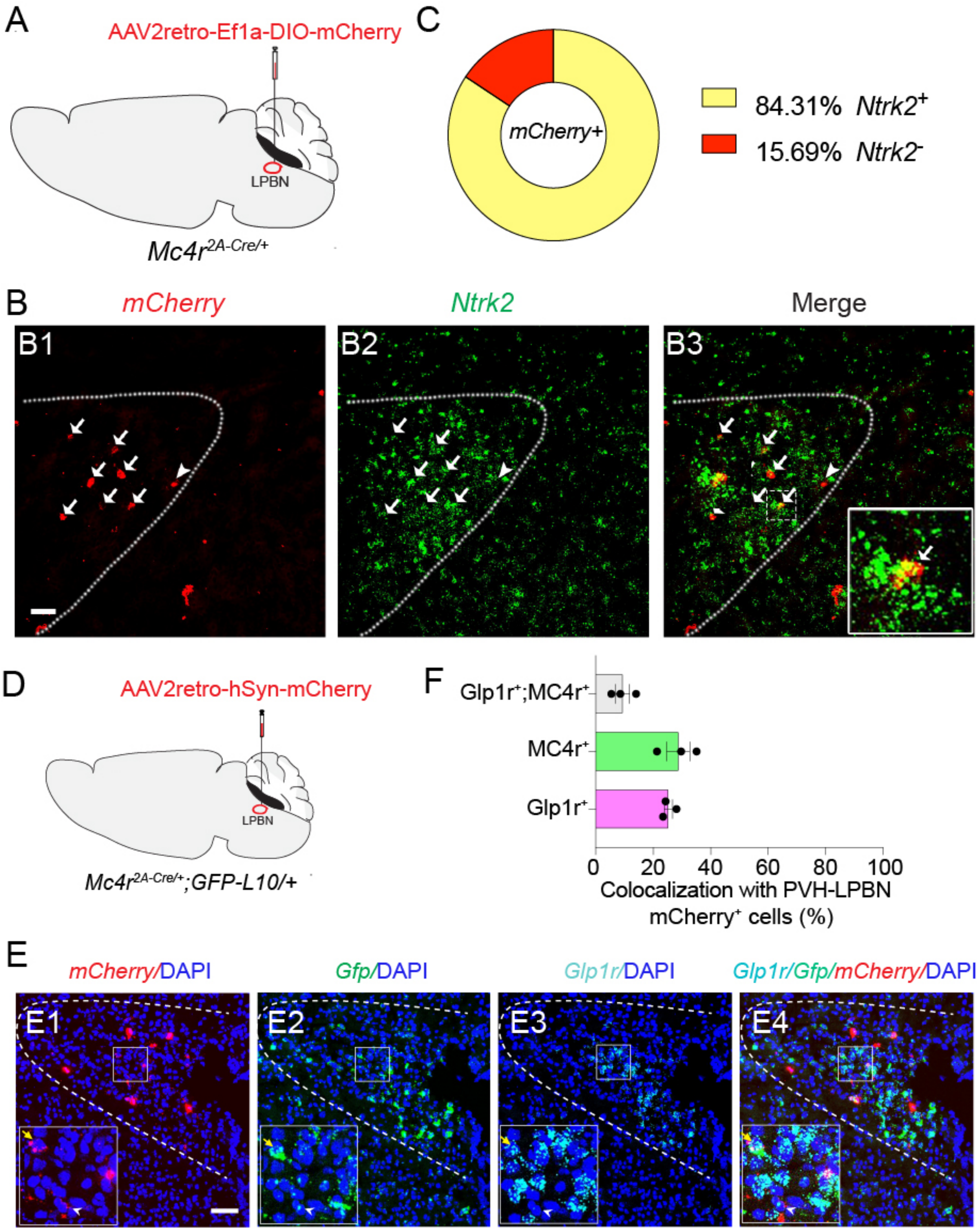


**Figure S1. Co-expression of *Ntrk2*, *Mc4r*, and *Glp1r* in the PVH.**

(A) Labeling LPBN-projecting PVH$^{MC4R}$ neurons with mCherry by injection of AAV2retro-Ef1a-DIO-mCherry into the LPBN of *Mc4r$^{2A\text{-}Cre/+}$* mice.

(B) Representative FISH micrographs showing co-localization of *mCherry* and *Ntrk2* mRNAs. Arrows denote cells co-expressing *mCherry* and *Ntrk2* mRNAs, whereas the arrowhead denotes

one cell that expresses *mCherry* mRNA but not *Ntrk2* mRNA. The insert in B3 shows a high-magnification image of the boxed area. Scale bar, 50 μm.
(C) Quantification of *mCherry* and *Ntrk2* mRNA colocalization. n=3 mice.
(D) Strategy for labeling LPBN-projecting PVH neurons with mCherry in *Mc4r*$^{2A\text{-}Cre/+}$;GFP-L10a/+ mice where MC4R cells express GFP.
(E) Representative FISH micrographs showing colocalization of *mCherry*, *Gfp* (indicative of *Mc4r* expression), and *Glp1r* mRNAs. Inserts show high-magnification images of the boxed areas. The yellow arrow denotes a cell expressing all three mRNAs, whereas the white arrowhead denotes a cell expressing *mCherry* and *Gfp* mRNAs but not *Glp1r* mRNA. Scale bar, 50 μm.
(F) Quantification of *Mc4r* and *Glp1r* mRNA expression in LPBN-projecting PVH neurons. n=3 mice.
All data are presented as mean ± SEM.

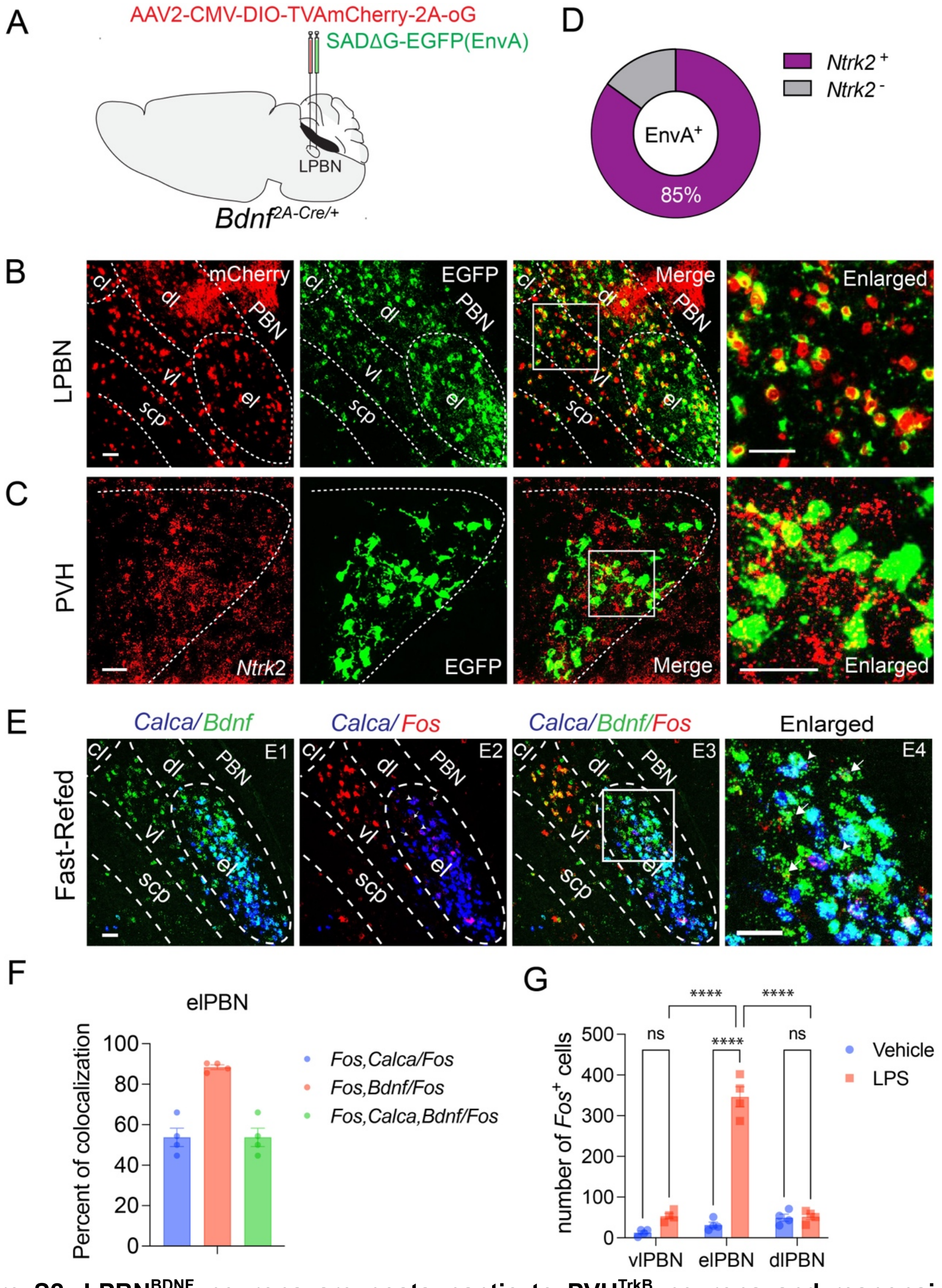


**Figure S2. LPBN$^{BDNF}$ neurons are postsynaptic to PVH$^{TrkB}$ neurons and responsive to refeeding.**

(A) Strategy for monosynaptic retrograde tracing of neurons presynaptic to LPBN$^{BDNF}$ neurons.

(B) Representative confocal micrographs showing LPBN$^{BDNF}$ neurons transduced by AAV2-CMV-DIO-TVAmCherry-2A-oG (expressing mCherry) and SADΔG-EGFP (EnvA) (expressing EGFP).

(C) Representative confocal micrographs showing that many EGFP-expressing neurons also express *Ntrk2* in the PVH.
(D) Quantification of *Ntrk2* and *Gfp* co-expression in the PVH. n=4 mice.
(E) Representative micrographs showing co-expression of *Calca*, *Bdnf*, and *Fos* mRNAs in the elPBN and vlPBN of mice refed for 2 hours following an overnight fast. In the enlarged image (E4), arrows denote cells expressing *Bdnf* and *Fos* mRNAs, whereas arrowheads denote cells expressing all three mRNAs.
(F) Quantification of co-expression between *Fos*, *Bdnf*, and *Calca* mRNAs in the elPBN of mice refed for 2 hours following an overnight fast. n=4 mice.
(G) Number of *Fos*$^+$ cells in the vlPBN, elPBN, and dlPBN of mice treated with either vehicle or LPS. n=4 mice. One-way ANOVA followed by multiple comparisons test, ns, not significant, ****$p<0.0001$.
All data are presented as mean ± SEM. Scale bars, 50 μm. Abbreviation: dl, dorsal lateral PBN; vl, ventral lateral PBN; el, external lateral PBN; scp, superior cerebellar peduncle.

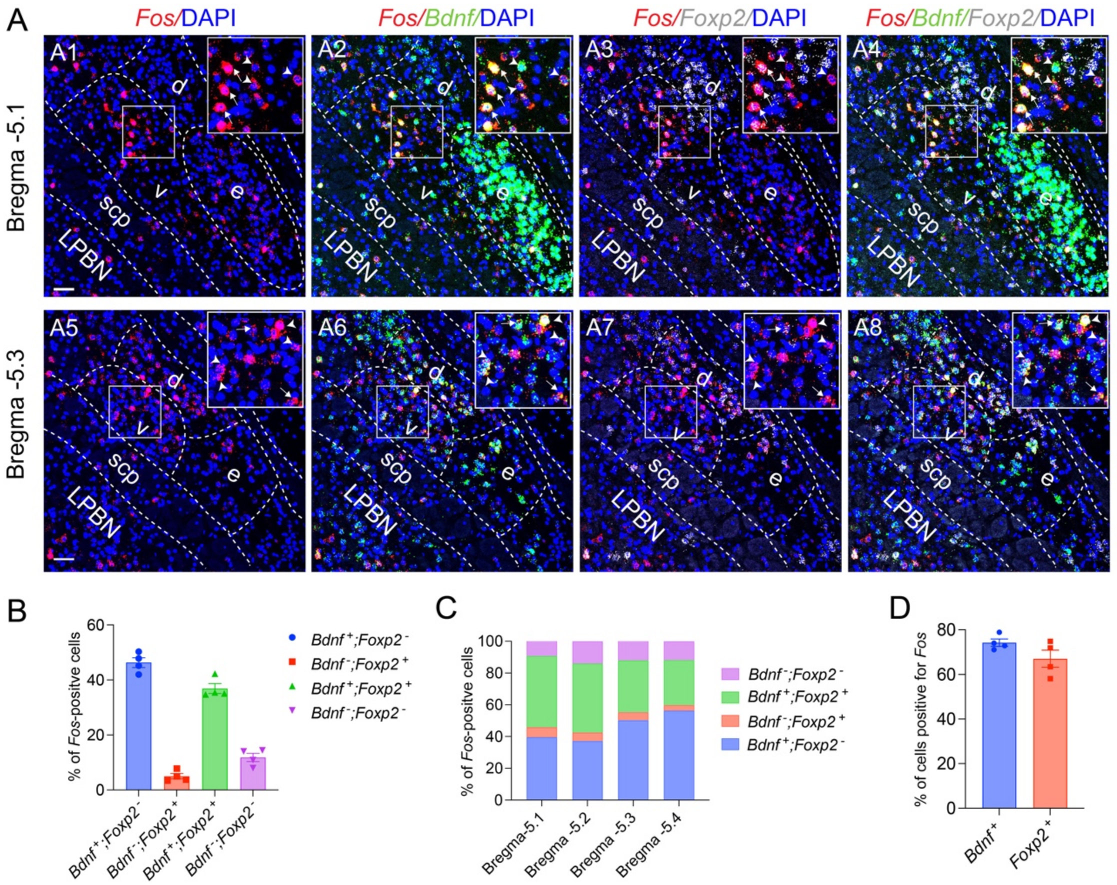


**Figure S3. Refeeding induced *Fos* expression in distinct groups of vlPBN neurons.**

(A) Representative FISH images showing *Fos*, *Bdnf*, and *Foxp2* mRNA expression at two rostrocaudal levels of the vlPBN in mice refed for 2 hours following an overnight fast. Scale bar, 50 μm. Abbreviations: d, dlPBN; e, elPBN; v, vlPBN; scp, superior cerebellar peduncle.

(B) Percentage of *Fos*-positive cells that express *Bdnf* and/or *Foxp2* in the vlPBN. n=4 mice.

(C) Percentage of *Fos*-positive vlPBN cells that co-express *Bdnf* mRNA only (blue), *Foxp2* mRNA only (orange), both transcripts (green), or neither transcript (purple) at four rostrocaudal levels of the vlPBN. n=4 mice.

(D) Percentage of *Bdnf* or *Foxp2* cells that are positive for *Fos*. n=4 mice.

All data are presented as mean ± SEM.

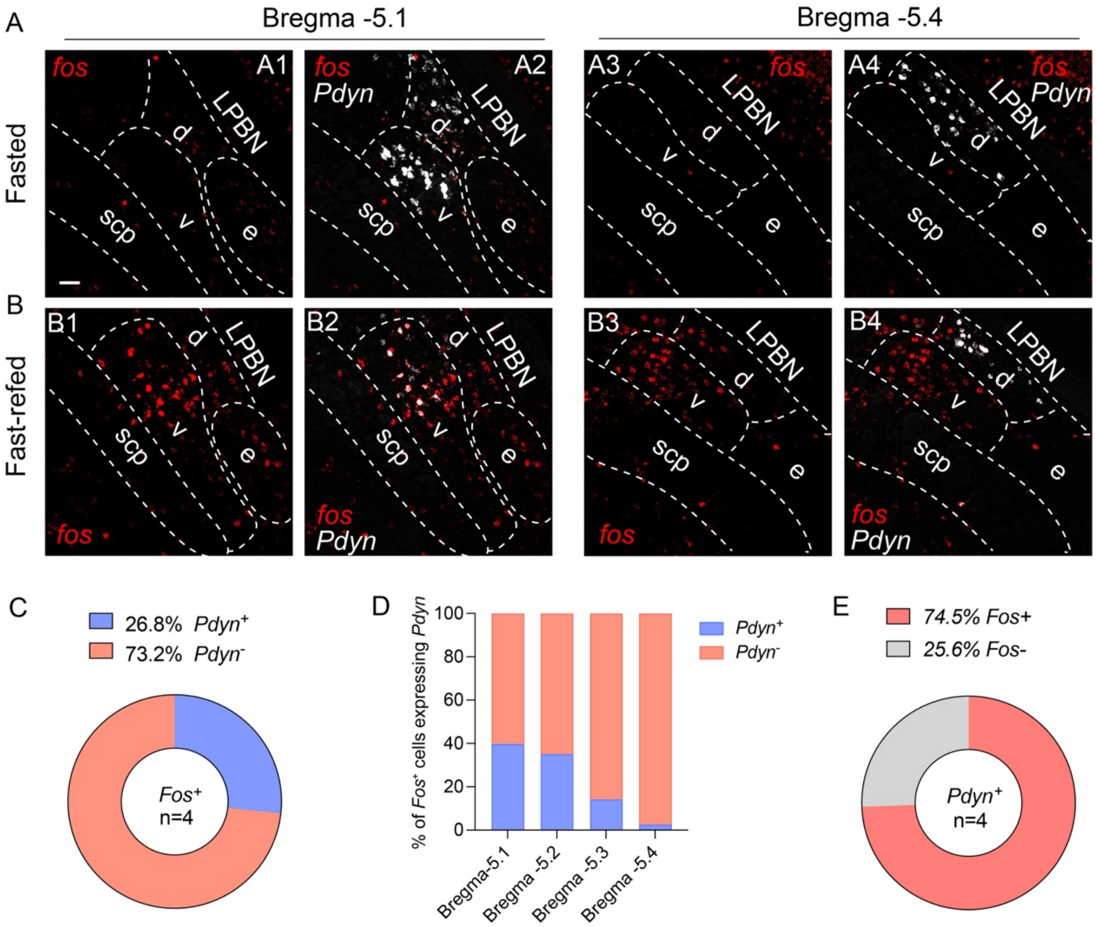


**Figure S4. Refeeding induced *Fos* expression in vlPBN$^{Pdyn}$ neurons.**

(A and B) Representative FISH images showing *Fos* and *Pdyn* mRNA expression at two rostrocaudal levels of the vlPBN in mice fasted overnight or refed for 2 hours following an overnight fast. Scale bar, 50 μm. Abbreviations: d, dlPBN; e, elPBN; v, vlPBN; scp, superior cerebellar peduncle.

(C) Proportion of *Fos*-positive vlPBN cells that expressed *Pdyn*. n=4 mice.

(D) Proportion of *Fos*-positive vlPBN neurons that either co-express *Pdyn* mRNA (blue) or lack *Pdyn* expression (orange) at four rostrocaudal levels of the vlPBN.

(E) Proportion of *Pdyn*-expressing vlPBN cells that were positive for *Fos*. n=4 mice.

All data are presented as mean ± SEM.

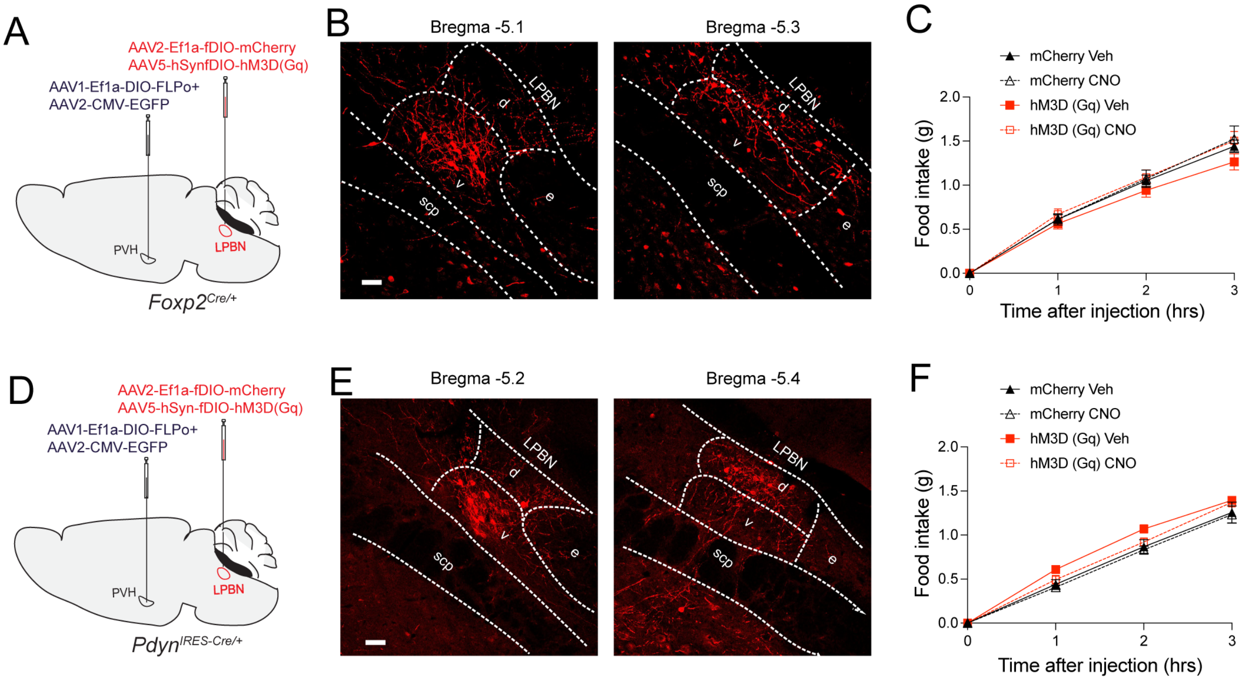


**Figure S5. Activation of PVH-innervated *Foxp2*- or *Pdyn*-expressing LPBN neurons did not affect food intake.**

(A) Strategy for selective expression of either mCherry or hM3D(Gq)-mCherry in PVH-innervated $\text{LPBN}^{\text{Foxp2}}$ neurons.

(B) mCherry expression at two rostrocaudal levels of the LPBN.

(C) Nocturnal food intake following vehicle (Veh) or CNO injection. n=6-7 mice per group. Two-way ANOVA followed by Tukey's multiple comparisons test, not significant.

(D) Strategy for selective expression of either mCherry or hM3D(Gq)-mCherry in PVH-innervated $\text{LPBN}^{\text{Pdyn}}$ neurons.

(E) mCherry expression at two rostrocaudal levels of the LPBN.

(F) Nocturnal food intake following vehicle (Veh) or CNO injection. n=6-7 mice per group. Two-way ANOVA followed by Tukey's multiple comparisons test, not significant.

All data are presented as mean ± SEM. Scale bars, 50 μm. Abbreviation: d, dorsal lateral PBN; e, external lateral PBN; v, ventral lateral PBN; scp, superior cerebellar peduncle.

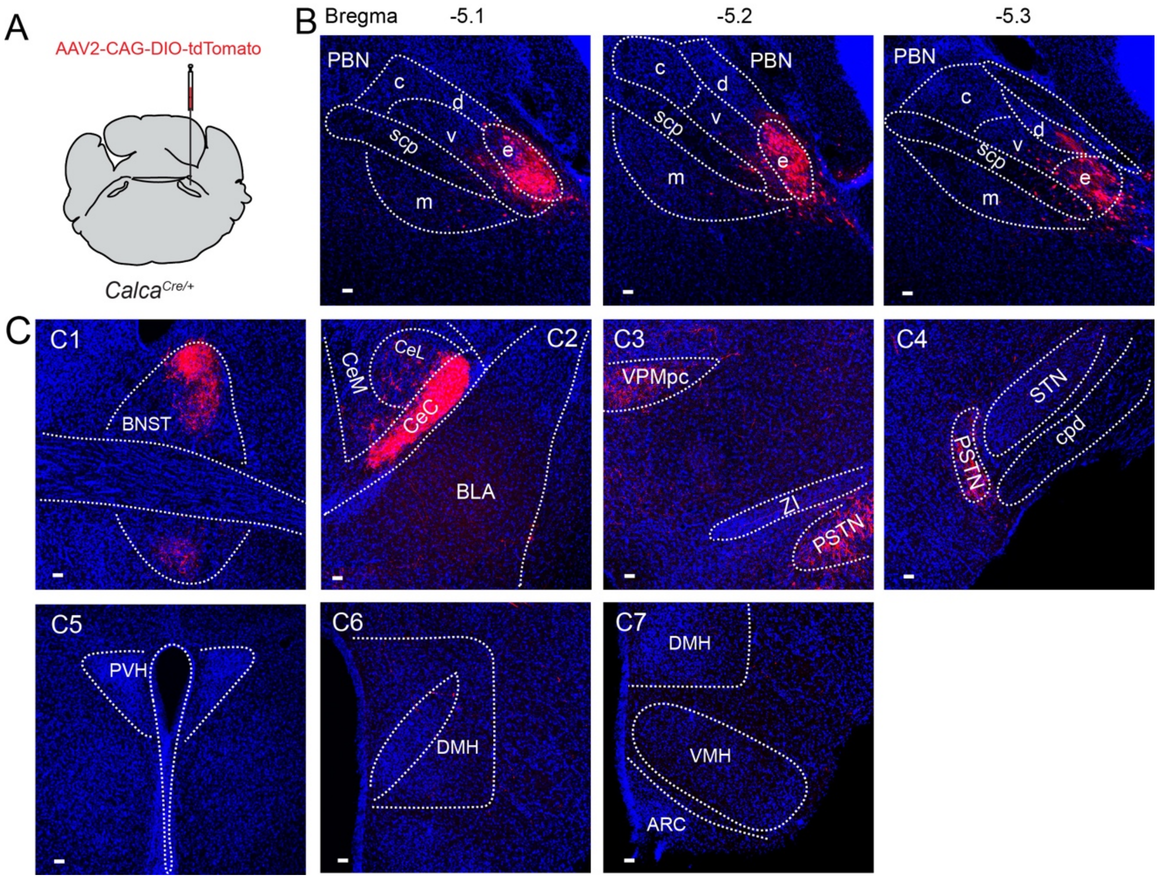


**Figure S6. Projections of elPBN$^{CGRP}$ neurons.**

(A) Strategy for selective labeling of elPBN$^{CGRP}$ neurons in *Calca*$^{Cre/+}$ mice with tdTomato.

(B) Labeled CGRP neurons in elPBN at three rostrocaudal levels. Scale bars, 50 μm.

(C) Axonal projections of elPBN$^{CGRP}$ neurons. Scale bars, 50 μm.

Abbreviations: c, central lateral PBN; d, dorsal lateral PBN; e, external lateral PBN; v, ventral lateral PBN; m, medial PBN; scp, superior cerebellar peduncle; BLA, basolateral amygdala; BNST, bed nucleus of the stria terminalis; CeC, CeL, and CeM, capsular, lateral, and medial division of the central amygdala, respectively; PSTN, parasubthalamic nucleus; STN, subthalamic nucleus; VPMpc, parvicellular part of the ventral posteromedial nucleus; ZI, zona incerta.

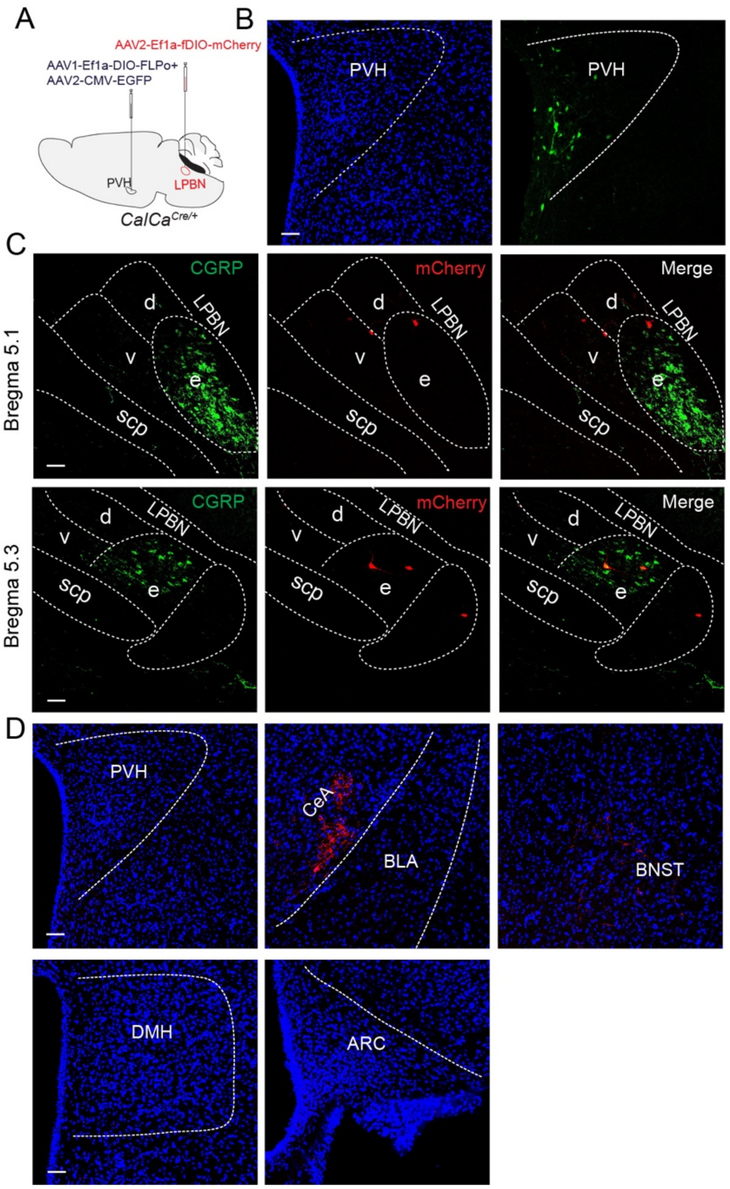


**Figure S7. Projections of PVH-innervated elPBN$^{CGRP}$ neurons.**

(A) Strategy for selective labeling of PVH-innervated elPBN$^{CGRP}$ neurons in *Calca*$^{Cre/+}$ mice with mCherry.

(B) The injection site marked by EGFP. Scale bar, 50 μm.

(C) mCherry-labeled CGRP neurons in the elPBN that is delineated by CGRP immunohistochemistry (green). Scale bar, 50 μm.

(D) mCherry-labeled axons from PVH-innervated elPBN$^{CGRP}$ neurons were detected in the CeA and BNST, whereas no axonal labeling was observed in the PVH, DMH, and ARC. Scale bars, 50 μm.

Abbreviations: d, dorsal lateral PBN; e, external lateral PBN; v, ventral lateral PBN; scp, superior cerebellar peduncle; ARC, arcuate nucleus; BLA, basolateral amygdala; BNST, bed nucleus of the stria terminalis; CeA, central amygdala; DMH, dorsomedial hypothalamus; PVH, paraventricular hypothalamus.

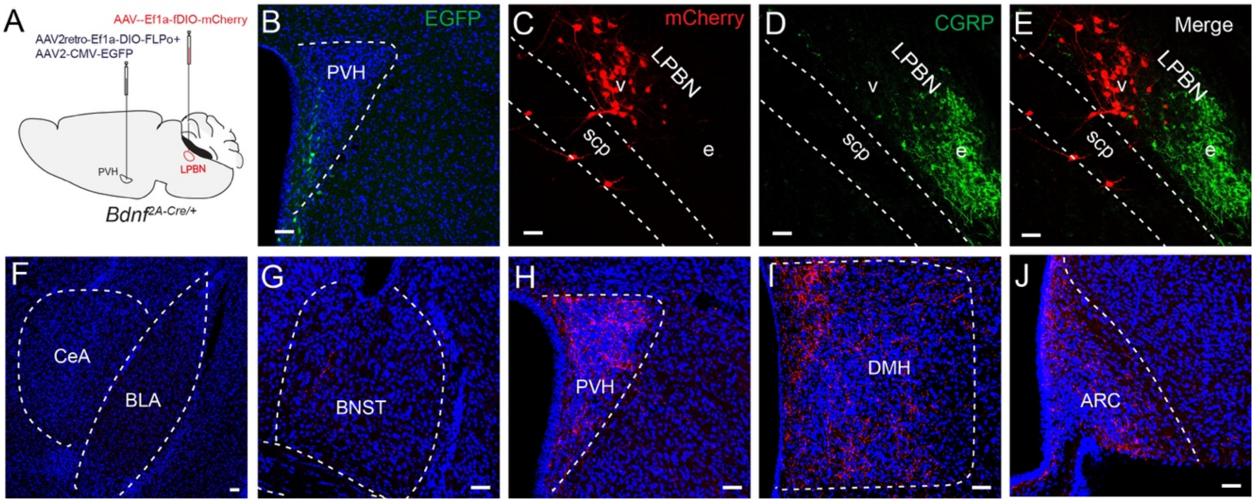


**Figure S8. Collaterals of PVH-projecting LPBN$^{BDNF}$ neurons.**

(A) Strategy for selective labeling of PVH-projecting LPBN$^{BDNF}$ neurons in *Bdnf*$^{2A\text{-}Cre/+}$ mice with mCherry.

(B) EGFP labeled the injection site in the PVH. Scale bar, 50 μm.

(C-E) mCherry-labeled PVH-projecting LPBN$^{BDNF}$ neurons. CGRP immunohistochemistry delineates the elPBN. Scale bars, 50 μm.

(F-J) mCherry-labeled axons from PVH-projecting LPBN$^{BDNF}$ neurons were detected in the PVH, DMH, and ARC (H-J), whereas little or no axonal labeling was observed in the CeA or BNST (F and G). Scale bars, 50 μm.

Abbreviations: e, external lateral PBN; v, ventral lateral PBN; m, medial PBN; scp, superior cerebellar peduncle; ARC, arcuate nucleus; BLA, basolateral amygdala; BNST, bed nucleus of the stria terminalis; CeA, central amygdala; DMH, dorsomedial hypothalamus; PVH, paraventricular hypothalamus.

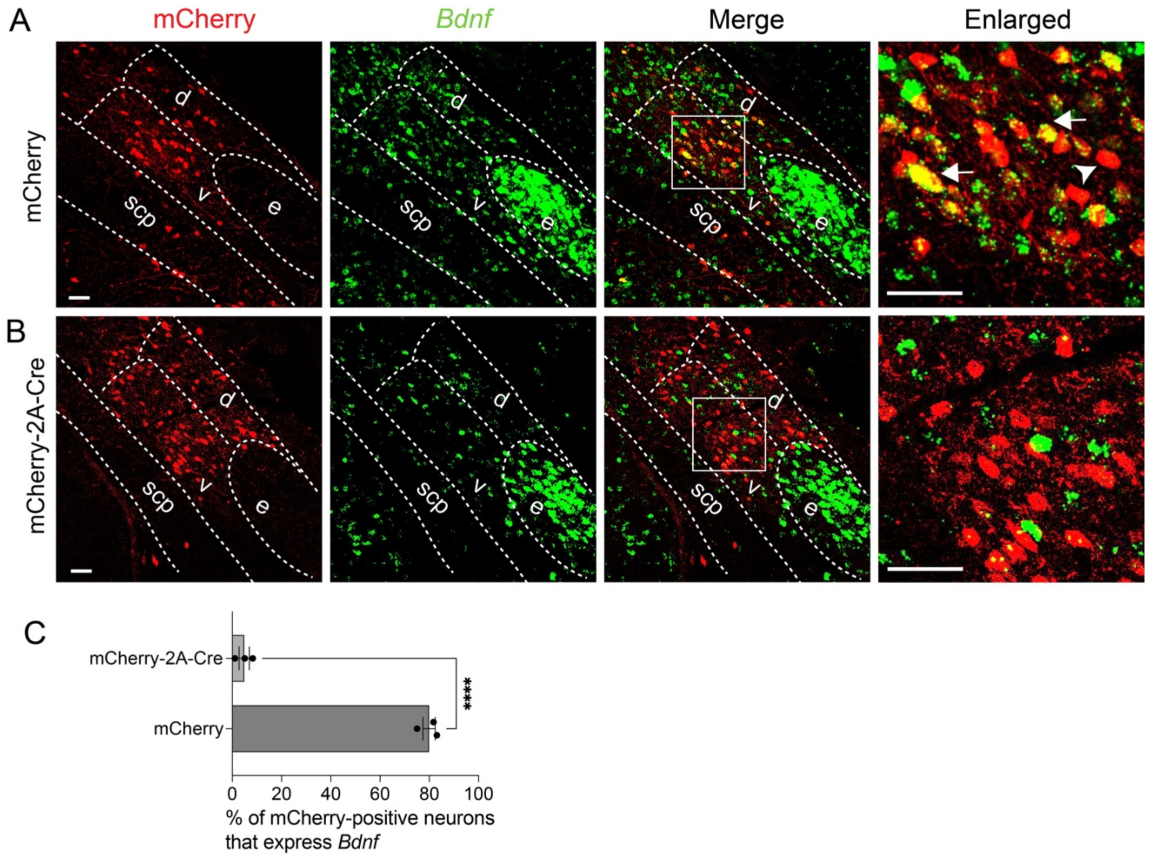


**Figure S9. *Bdnf* deletion in the vlPBN and dlPBN of *Bdnf*$^{lox/lox}$ mice.**

(A) mCherry immunohistochemistry and *Bdnf* mRNA FISH in the LPBN of *Bdnf*$^{lox/lox}$ mice injected with AAV2-Ef1a-fDIO-mCherry. Scale bars, 50 μm.

(B) mCherry immunohistochemistry and *Bdnf* mRNA FISH in the LPBN of *Bdnf*$^{lox/lox}$ mice injected with AAV2-Ef1a-fDIO-mCherry-2A-Cre. Scale bars, 50 μm.

(C) Quantification of mCherry-positive neurons in the vlPBN that expressed *Bdnf*. An mCherry-positive neuron containing 2 or more *Bdnf* FISH puncta was considered a *Bdnf*-expressing cell. Unpaired t test, ****p<0.0001; n=3 mice per group.

Data are presented as mean ± SEM. Abbreviations: d, dorsal lateral PBN; e, external lateral PBN; v, ventral lateral PBN; scp, superior cerebellar peduncle.
